# Oxygen-generating cryogels restore T cell-mediated cytotoxicity in hypoxic tumors

**DOI:** 10.1101/2020.10.08.329805

**Authors:** Thibault Colombani, Loek J. Eggermont, Stephen M. Hatfield, Mahboobeh Rezaeeyazdi, Adnan Memic, Michail V. Sitkovsky, Sidi A. Bencherif

**Affiliations:** Department of Chemical Engineering, Northeastern University, Boston, MA 02115, USA; New England Inflammation and Tissue Protection Institute, Northeastern University, 360 Huntington Avenue, Boston, MA 02115, USA; Center of Nanotechnology, King Abdulaziz University, Jeddah, Makkah 21589, Saudi Arabia; Department of Bioengineering, Northeastern University, Boston, MA 02115, USA; Harvard John A. Paulson School of Engineering and Applied Sciences, Harvard University, Cambridge, MA 02138, USA; Biomechanics and Bioengineering (BMBI), UTC CNRS UMR 7338, University of Technology of Compiègne, Sorbonne University, 60203 Compiègne, France

**Keywords:** Cryogels, Hypoxia, Cancer, Oxygenation, T cell-mediated Antitumor Activity

## Abstract

Solid tumors are protected from antitumor immune responses due to their hypoxic microenvironments. Weakening hypoxia-driven immunosuppression by hyperoxic breathing of 60% oxygen has shown to be effective in unleashing antitumor immune cells against solid tumors. However, efficacy of systemic oxygenation is limited against solid tumors outside of lungs. Therefore, it is essential to develop targeted oxygenation alternatives to weaken tumor hypoxia as novel approaches to cancer immunotherapies. Herein, we report on injectable oxygen-generating cryogels (O2-cryogels) to reverse tumor-induced hypoxia. These macroporous biomaterials were designed to locally deliver oxygen, inhibit the expression of hypoxia-inducible genes in hypoxic melanoma cells, and reduce the accumulation of immunosuppressive extracellular adenosine. O2-cryogels enhance T cell-mediated secretion of cytotoxic proteins, restoring the killing ability of tumor-specific CTLs, both in vitro and in vivo. In summary, O2-cryogels provide a unique and safe platform to supply oxygen as a co-adjuvant in hypoxic tumors and improve cancer immunotherapies.

## Introduction

Over the last decade, immunotherapy has evolved into a very promising new frontier for fighting various types of cancer.(*1*) A number of these immunotherapies are either approved for use or are under clinical trials, particularly immune checkpoint inhibitors (e.g., anti-PD-1, anti-PD-L1, and/or anti-CTLA-4 antibodies) and T cell therapy (e.g., chimeric antigen receptor [CAR] T cells).(*2-4*) However, their limited success in solid tumors has been attributed to tumor-protecting mechanisms that lead to immune escape.(*5*) Low oxygen tension (< 3% O_2_), or hypoxia, is a common feature of solid tumors.(*6*) Hypoxia within the tumor microenvironment (TME) is a major impediment to effective cancer therapies and has been associated with poor clinical outcomes.(*7*) Hypoxic tumors often exhibit poor responses to systemic therapies(*8*) and radiotherapy,(*9, 10*) resistance to apoptosis,(*11*) and an aggressive phenotype that leads to rapid disease progression and poor prognosis.(*12, 13*)

Hypoxic stress is a key factor involved in creating an immunosuppressive environment and as a result, inhibits incoming cytotoxic T lymphocytes (CTLs).(*14-17*) These immunosuppressive effects are primarily mediated by hypoxia-inducible factors (HIFs)(*18-20*) and A2A adenosine receptors (A2ARs).(*21, 22*) Tumor hypoxia stabilizes the transcription factor HIF1α,(*23*) leading to increased expression of the adenosine-generating ecto-enzymes CD39 and CD73.(*24*) This results in the accumulation of extracellular adenosine in the TME, which signals through cAMP-elevating A2ARs on the surface of immune cells.(*25-27*) Adenosine signaling suppresses the antitumor activity of T cells and natural killer (NK) cells, particularly their cytotoxic effector functions and secretion of important proinflammatory cytokines such as interferon gamma (IFNγ).(*28, 29*)

Several classes of therapeutics can reverse hypoxia-induced immunosuppression,(*30*) including A2AR antagonists,(*31, 32*) HIF inhibitors,(*30, 33*) and supplemental oxygenation therapy.(*24, 34*) Recent clinical studies using A2AR antagonists have shown encouraging results in targeting the hypoxia-adenosinergic pathway.(*35*) Similarly, systemic oxygen therapy such as respiratory hyperoxia has proven effective to enhance antitumor immunity in preclinical studies.(*22*) However, respiratory hyperoxia (> 60% O_2_) has been associated with inflammatory changes, alveolar infiltration, and, eventually, pulmonary fibrosis.(*36*) Furthermore, although safer, respiratory hyperoxia with lower oxygen concentrations (≤ 60% O_2_) is less effective against solid tumors in tissues distant from the lungs and it may exacerbate inflammatory events in patients.(*22, 36*) Therefore, there is a critical need to develop new strategies to target oxygen delivery and minimize off-target toxicities. Such approaches would be useful to reduce or eliminate local tumor hypoxia, create an immunopermissive TME, and ultimately enhance the effectiveness of cancer treatments.

Over the past decade, innovative biomaterials have gained great momentum as a means to study or modulate biological environments.(*37, 38*) Recent studies have shown that incorporating peroxides, percarbonate, or hydrogen peroxide (H_2_O_2_)-based particles in biomaterials allowed oxygen delivery within tissues.(*39, 40*) These biomaterials release oxygen in water(*41, 42*) and are typically coupled with an antioxidant to attenuate harmful free radical byproducts.(*39, 40, 43*) However, the administration of biomaterials is often invasive, which limits their clinical feasibility.(*44-46*) Furthermore, biomaterials that release oxygen require a highly interconnected macroporous network to tackle mass transport and diffusion limitations,(*47*) but such structures are challenging to engineer.^(*48*)^ Recently, cryogel scaffolds have emerged as promising biomaterials that can be injected through conventional small-bore needles for minimally invasive delivery.(*49, 50*) This new class of injectable hydrogels possesses unique features such as shape-memory properties, mechanical flexibility, and a highly interconnected macroporous architecture essential for oxygen diffusion and availability throughout the construct.

We hypothesized that an oxygen-generating biomaterial capable of reversing tumor-induced hypoxia may reactivate inhibited T-cell function and activity, and subsequently reinforce T cell-mediated cytolytic activity against tumor cells. In this work, we engineered injectable macroporous oxygen-generating cryogels (O_2_-cryogels) that can reoxygenate O_2_-deprived TME, reverse hypoxia-driven immunosuppression, and ultimately restore antitumor CTL activity. The cryogels were fabricated with the naturally occurring glycosaminoglycan hyaluronic acid (HA), one of the major components of the tumor extracellular matrix (ECM). The three-dimensional (3D) cryogels were hybridized with calcium peroxide (CaO_2_) microparticles and conjugated to catalase (CAT) to boost local oxygen release while minimizing cytotoxicity. We first characterized the physical properties of O_2_-cryogels, including their unique interconnected macroporous networks, shape memory properties, and syringe injectability. Next, we investigated the ability of O_2_-cryogels to release oxygen in a controlled and sustained fashion under normoxic (20% O_2_) and hypoxic (1% O_2_) conditions while being inherently cytocompatible and biocompatible. We also studied the potential of O_2_-cryogels to reverse tumor hypoxia, downregulate expression levels of hypoxia-induced proteins, increase proinflammatory cytokines, and decrease immunosuppressive molecules. Finally, in comparison to respiratory hyperoxia (60% O_2_), we evaluated the ability of O_2_-cryogels to restore the antitumor activity of inhibited hypoxic T cells in hypoxia and their use as an immunological co-adjuvant in a highly aggressive and advanced-stage murine melanoma model. This study reports on (*i*) a strategy to re-establish T cell function when subjected to hypoxic stress and (*ii*) the potential of biomaterial-mediated intratumoral oxygen delivery as a safe co-adjuvant to reinforce pre-existing antitumoral immune response in vivo.

## Results

### Fabrication of O_2_-cryogels

O_2_-cryogels were fabricated by free radical polymerization of HAGM, APR, and APC at −20 °C in the presence of various concentrations of CaO_2_ (Figure 1a). HAGM was incorporated into the cryogel backbone to mimic aspects of the TME,(*51*) APR to prevent anoikis and promote cell–cryogel interactions,(*52*) and CaO_2_ and APC to release oxygen and deplete H_2_O_2_ byproduct, respectively. During cryopolymerization, the macromonomers, CaO_2_, and initiators are phase-separated from ice crystals (step 1), enabling polymerization within the unfrozen liquid phase and CaO_2_ particle entrapment in the newly formed polymer walls (step 2). Once the reaction is complete, the cryogels are thawed at RT to melt the ice crystals, leaving behind a polymer network with large and interconnected pores (step 3).

**Figure 1.**
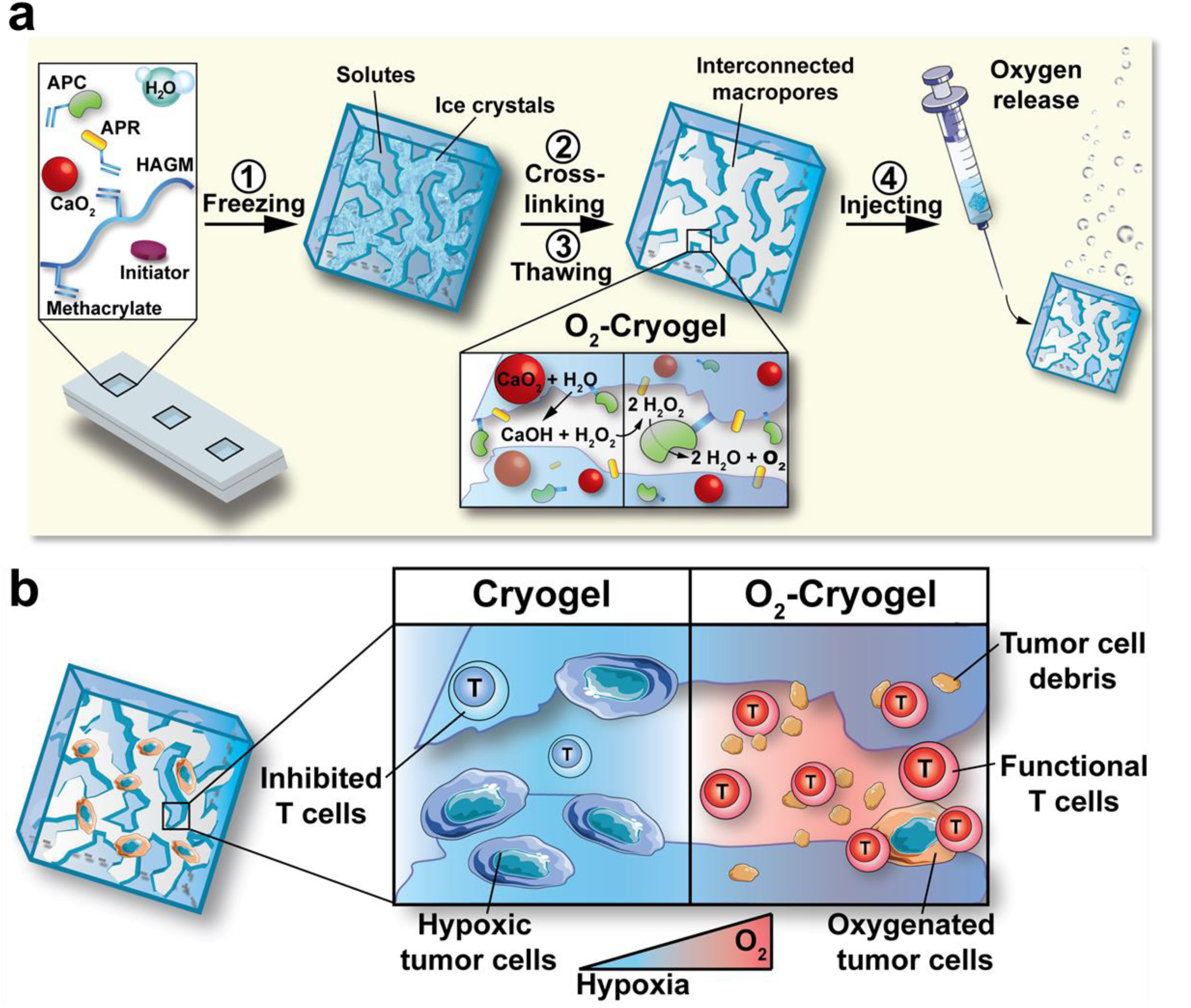
Schematic describing the fabrication process of O_2_-cryogels and their ability to restore T cell-mediated cytotoxicity in a hypoxic tumor microenvironment. **a**) To fabricate O_2_-cryogels, hyaluronic acid grafted with methacrylate residues (HAGM), calcium peroxide CaO_2_ particles, acrylate-PEG-catalase (APC), acrylate-PEG_3500_-G_4_RGDSP (APR), and ammonium persulfate/ tetramethylethylenediamine (initiator system) were mixed and incubated at −20 °C for 16 h to allow cryopolymerization. Steps 1,2: This process involves a phase separation with ice crystal formation, solutes concentration (polymer, APC, APR, CaO_2_), and crosslinking within a non-frozen liquid phase leading to CaO_2_ particle entrapment within the dense polymer walls. Step 3: A thawing process was performed to remove ice crystals and leave an open interconnected macroporous network. Step 4: The mechanically robust O_2_-cryogels sustained syringe injection using a conventional small-bore needle and released oxygen in a controlled manner when hydrated. The schematic shows the active release of oxygen bubbles by O_2_-cryogels in water. **b**) In hypoxia, tumor cell-laden cryogels can generate an immunosuppressive environment that inhibits cytotoxic T cells and prevents their cytotoxic function (left panel). Oxygen-releasing O_2_-cryogels can reverse local hypoxia and restore the ability of T cells to target tumor cells, T cell-tumor cell interactions, and the secretion of cytotoxic proteins (e.g., perforin and granzyme B), leading to apoptotic tumor cell death and tumor cell debris (right panel).

Unlike other large-scale oxygen-generating biomaterials, O_2_-cryogels were designed to withstand substantial mechanical deformation and compression, resulting in a minimally invasive biomaterial that can overcome the shear stress experienced during syringe injection (step 4). O_2_-cryogels were engineered to controllably release oxygen via CaO_2_ decomposition in water. We hypothesized that local oxygen delivery would relieve the hypoxic stress and as a result, (1) reverse the immunosuppressive TME and (2) restore impaired effector T-cell function. This strategy is intended to facilitate CTLs to infiltrate solid tumors and ultimately kill tumor cells (Figure 1b).

### Characterization of O_2_-cryogels

The controlled and sustained release of oxygen from O_2_-cryogels hinges on the efficient encapsulation and distribution of CaO_2_ particles throughout the polymer network. To demonstrate the successful and homogeneous incorporation of the particles into the polymer walls, cryogels and O_2_-cryogels containing different amounts of CaO_2_ (0.1–1% wt/vol) were stained with Alizarin Red S before and after syringe injection through a 16-gauge needle (Figure 2a). The red staining intensity increased with increasing CaO_2_ concentration, confirming the efficient incorporation of these particles within O_2_-cryogels. Furthermore, O_2_-cryogels exhibited the same shape-memory properties as those of their cryogel analogues (Supplementary Video 1), and no significant changes in Alizarin Red S staining were observed following injection. Confocal microscopy (Figure 2b–g) combined with scanning electron microscopy imaging (Figure 2 h–i and Supplementary Figure 1) and EDX analysis (Figure 2j–k and Supplementary Figure 2) validated the Alizarin Red S staining results. Specifically, the particle entrapment within O_2_-cryogel polymer walls increased proportionally to the initial CaO_2_ loading. Additionally, the particles were homogeneously distributed throughout the O_2_-cryogel construct. Altogether, these results suggest that CaO_2_ particles remain physically entrapped within polymer walls even under an applied shear stress.

**Figure 2.**
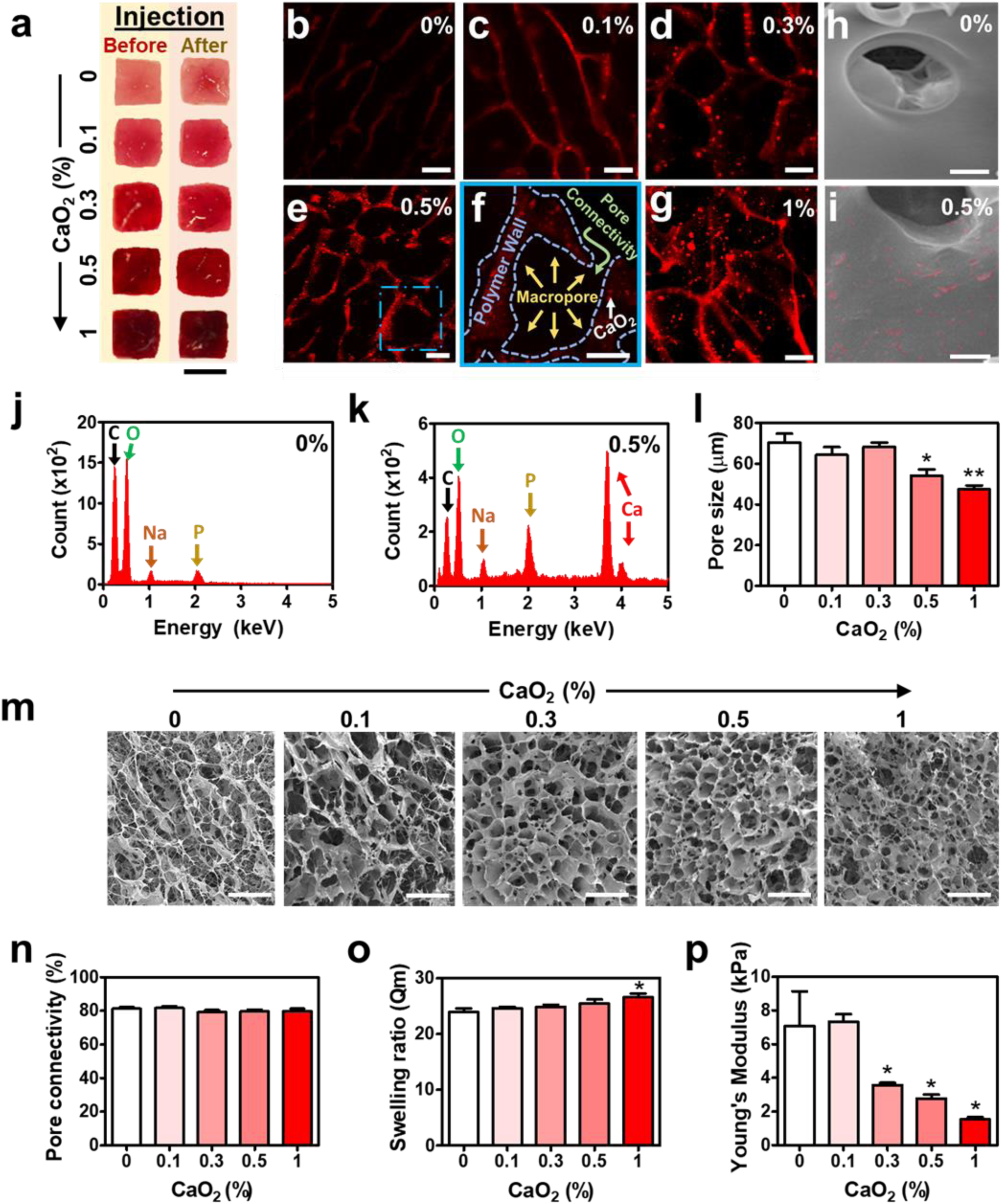
Physical characteristics of O_2_-cryogels. O_2_-cryogels were prepared with 0.1, 0.3, 0.5, or 1% wt/v CaO_2_ and their physical features compared to cryogels. **a–i**) Entrapped calcium peroxide particles within the O_2_-cryogel polymer walls. a) Photographs showing cryogels and O_2_-cryogels stained with Alizarin Red S before and after syringe injection. b–g) Confocal images showing CaO_2_ particles (stained with Alizarin Red S) physically entrapped within O_2_-cryogel polymer walls at various concentrations (c-g) and compared to cryogels (b). The confocal image (f) highlights the hierarchical macroporous structure of O_2_-cryogels containing 0.5% CaO_2_ (entrapped CaO_2_ particles, crosslinked polymer walls, and interconnected macropores). h,i) Scanning electron microscopy images showing cryogels (h) and entrapped particles within the polymer walls of O_2_-cryogels containing 0.5% CaO_2_ (i). Pseudocoloring in pink is used to highlight CaO_2_. **j, k**) Elemental analysis of cryogels and O_2_-cryogels containing 0.5% CaO_2_. **l**) Pore size of cryogels and O_2_-cryogels. **m**) Scanning electron microscopy images showing the interconnected macroporous structure of cryogels and O_2_-cryogels. **n–p**) Physical characterization. n) Pore connectivity, **o**) Mass swelling ratios, and **p**) Young’s modulus values of cryogels and O_2_-cryogels. Each image or photograph is representative of n = 5 samples. Values represent the mean ± SEM (n = 6 cryogels per condition). Data were analyzed using ANOVA and Dunnett’s post-hoc test (compared to cryogels); ^*^P < 0.05, ^**^P < 0.01. Scale bars = 4 mm (a), 50 µm (b-g), 25 µm (i), and 200 µm (m).

Cryogels consist of a highly interconnected polymer network.^49^ However, the impact of CaO_2_ encapsulation on the physical properties of cryogels has never been investigated. Therefore, we evaluated the change in the cryogel macrostructure as a function of CaO_2_ concentration, using scanning electron microscopy (Figure 2l–m). O_2_-cryogels displayed large pores ranging from 20 to 100 µm in diameter, and the pore size decreased proportionally with increasing CaO_2_ concentration—from 70.51 ± 10.7 µm (0% CaO_2_) to 47.43 ± 6.7 µm (1% CaO_2_). Next, the impact of the network microstructure of O_2_-cryogels on mechanical properties was assessed (Figure 2n–p). The addition of CaO_2_ did not change their pore connectivity (80 ± 3%) (Figure 2n). However, an increase in CaO_2_ concentration slightly enhanced their swelling ratio, from 23.9 ± 0.9 (0% CaO_2_) to 26.6 ± 0.6 (1% CaO_2_) (Figure 2o). Additionally, CaO_2_ encapsulation dramatically affected the Young’s modulus of the cryogels during compression (Figure 2p). While cryogels displayed a modulus of 7 ± 2 kPa, an increase in CaO_2_ concentration to 1% CaO_2_ reduced the Young’s modulus value down to 1.5 ± 0.2 kPa. Collectively, these results indicate that CaO_2_ encapsulation slightly impacted the physical properties of the cryogels without altering their unique interconnected macroporous structure, injectability, and shape-memory properties.

### Controlled and sustained release of oxygen from O_2_-cryogels

In water, CaO_2_ slowly decomposes to form calcium hydroxide (Ca(OH)_2_) and oxygen as depicted in Equations 2 and 3:

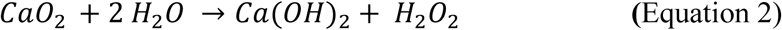

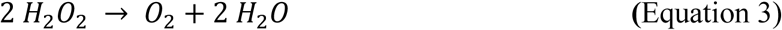

However, this reaction also produces the reactive oxygen species H_2_O_2_ as a byproduct, which can be harmful to cells, especially at concentrations beyond 10 µM.(*53*) To prevent H_2_O_2_-related cytotoxicity, CAT was chemically modified with ACRL-PEG-NHS to form APC. Next, APC was covalently incorporated into O_2_-cryogel during cryopolymerization to sustainably degrade H_2_O_2_ into H_2_O and O_2_. Compared to non-modified CAT, the enzymatic activity of modified CAT (i.e., APC) was decreased by 70 ± 3% (Supplementary Figure 3a). Next, the ability of APC to deplete H_2_O_2_ following cryogelation was investigated. APC-containing or APC-free O_2_-cryogels were incubated in water at 37 °C for 15 min, after which the production of H_2_O_2_ was evaluated. Incorporating APC in O_2_-cryogels containing 0.1% to 0.3% CaO_2_ effectively prevented any residual H_2_O_2_ release (Figure 3a). Similarly, APC-grafted O_2_-cryogels containing 0.5% and 1% CaO_2_ achieved low H_2_O_2_ levels of 4 ± 1 and 11 ± 2.5 µM, respectively. APC also promoted oxygen production from H_2_O_2_ (Supplementary Figure 3b). While no oxygen release occurred when APC-free cryogels were immersed in 50 mM H_2_O_2_ (Supplementary Video 2), cryogels containing 1% APC produced small gas bubbles under the same conditions, suggesting rapid conversion of H_2_O_2_ to oxygen (Supplementary Video 3).

**Figure 3.**
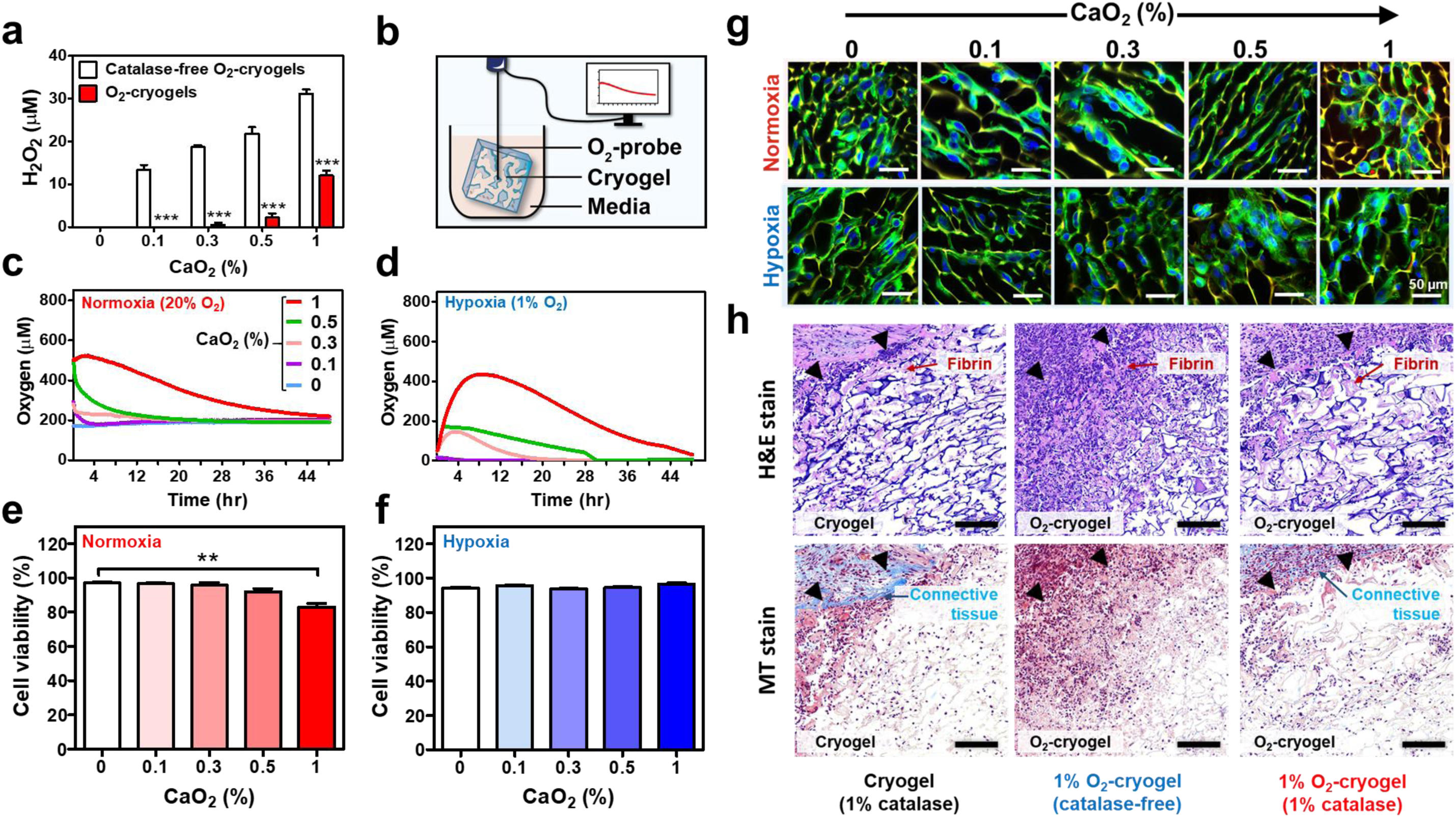
O_2_-cryogels enable controlled and sustained release of oxygen and are cytocompatible (in vitro) and biocompatible (in vivo). **a**) H_2_O_2_ release from catalase-free (white) and (red, catalase-containing) O_2_-cryogels. **b**) Oxygen measurement setup. Cryogels and O_2_-cryogels were individually immersed in 2-mL Eppendorf tubes containing cell culture medium and incubated at 37 °C under normoxic (20% O_2_) or hypoxic (1% O_2_) conditions. A needle-type optical oxygen microprobe was inserted in the center of each cryogel, and changes in the dissolved oxygen concentration were recorded during the course of the experiment. **c, d**) Controlled oxygen release from cryogels and O_2_-cryogels under normoxic (c) and hypoxic conditions (d). **e, f**) B16-F10 cell viability after 24 h of incubation within cryogels and O_2_-cryogels under (e) normoxic (20% O_2_) and (f) hypoxic (1% O_2_) conditions. **g**) Representative confocal images of B16-F10 cell viability after 24 h of incubation within cryogels and O_2_-cryogels under normoxic (top) and hypoxic (bottom) conditions. Blue = nuclei stained with DAPI, red = dead cells stained with ViaQuant Far Red, green = actin cytoskeleton stained with Alexa Fluor 488 phalloidin, yellow = polymer walls stained with rhodamine. **h**) Histological analysis of explanted cryogels (cryogel) containing 1% catalase, O_2_-cryogels containing 1% CaO_2_ (1% O_2_-cryogel) and 1% catalase, and catalase-free 1% O_2_-cryogels on day 7 post-subcutaneous injection using H&E and MT staining. The arrows indicate the boundary between the cryogel and host tissue. Confocal and histological images are representative of n = 5 samples per condition. Values represent the mean ± SEM (n = 6 cryogels). Data in panels (a, e, f) were analyzed using ANOVA and Dunnett’s post-hoc test (compared to catalase-free O_2_-cryogels); ^*^P < 0.05, ^**^P < 0.01, ^***^P < 0.001. Scale bars = 50 µm (g) and 200 µm (h).

Next, we characterized the kinetics of oxygen release from cryogels containing varying amounts of CaO_2_ (0 to 1% wt/vol) by measuring the concentration of dissolved oxygen in the medium at 37 °C every 5 min for two days (Figure 3b and Supplementary Figure 3c). The amount of oxygen generated by O_2_-cryogels was proportional to the CaO_2_ content under normoxic (Figure 3c) and hypoxic conditions (Figure 3d). In normoxic conditions, O_2_-cryogels induced a maximum oxygen concentration increase in the medium (up to 215%) and achieved a sustained release of oxygen for up to 48 h (Figure 3c). As expected, cryogels did not release any oxygen. Similar results were also observed with preconditioned hypoxic (1% O_2_) media (Figure 3d). O_2_-cryogels increased the oxygen concentration in the medium up to 2000% of the initial concentration and allowed a sustained oxygen release for up to 48 h. In hypoxia, O_2_-cryogels released effectively oxygen and were able to reach oxygen levels above physioxia (5–8% O_2_, i.e., 80 μM of O_2_ at 37 °C, 1013 hPa) for 8 h (with 0.3% CaO_2_) and 40 h (with 1% CaO_2_).

These O_2_-cryogels were designed to be injectable. Therefore, their capacity to efficiently release oxygen upon injection was also evaluated. Because our experimental design required large cryogel cylinders, each O_2_-cryogel was compressed by up to 90% of their volume and subjected to shear stress to mimic the injection process (Supplementary Figure 4). Regardless of the CaO_2_ concentration, the cryogels showed the same oxygen release kinetics before and after compression, consistent with the CaO_2_ entrapment measurements (Figure 2a). Collectively, these results indicate that injectable oxygen-releasing cryogels were successfully fabricated and can sustainably deliver oxygen for up to 48 h under hypoxic and normoxic conditions.

**Figure 4.**
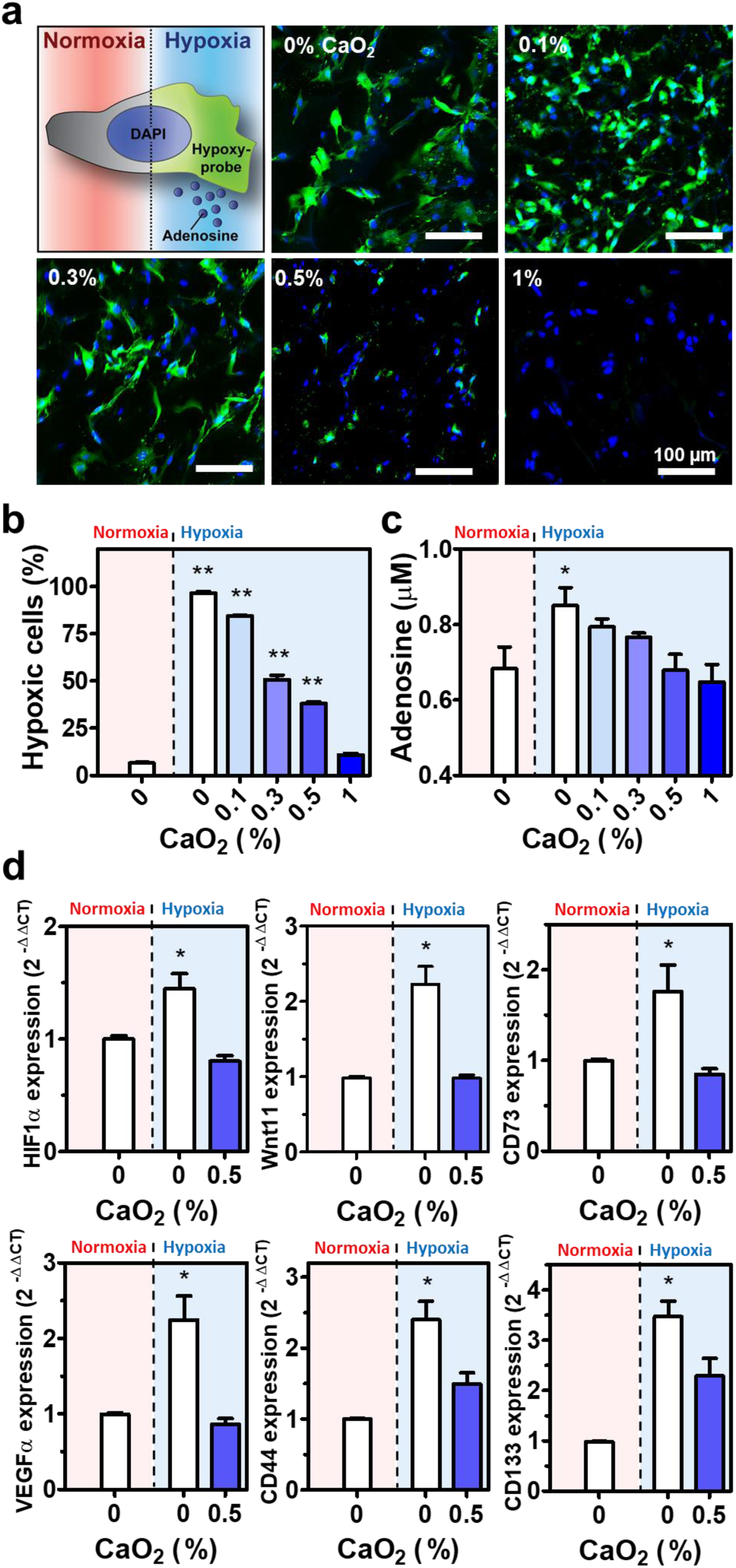
Cellular hypoxia reversal in melanoma cells via O_2_-cryogel-mediated oxygenation. **a**) Detection of hypoxia in B16-F10 cells in cryogels and O_2_-cryogels cultured in 1% O_2_ for 24 h. Blue staining = nuclei (stained with DAPI), green = hypoxic cells stained with Hypoxyprobe™-1. The data are representative of n = 4 samples per condition. **b**) Quantification of cellular hypoxia following a 24-h incubation period in cryogels and O_2_-cryogels under normoxic or hypoxic conditions. **c**) Adenosine concentration in B16-F10 cell supernatants after 24 h of incubation in cryogels and O_2_-cryogels under normoxic and hypoxic conditions. **d**) HIF1α, Wnt11, CD73, VEGFα, CD44, and CD133 expression levels in B16-F10 cells after 24 h of incubation in cryogels and O_2_-cryogels containing 0.5% CaO_2_ under normoxic and hypoxic conditions. Gene expression was determined by RT-qPCR, normalized against the expression level of HPRT (housekeeping gene) and compared to that in cell-laden cryogels in normoxia (2^-ΔΔCT^ = 1). Values represent the mean ± SEM (n = 3**–**5). Data were analyzed using ANOVA and Dunnett’s post-hoc test (compared to cryogels in normoxia); ^*^P < 0.05, ^**^P < 0.01. Scale bar = 100 µm.

### Cytocompatibility (in vitro) and biocompatibility (in vivo) of O_2_-cryogels

We next examined cell viability within O_2_-cryogels as an indication of cytocompatibility. Cryogels and O_2_-cryogels were partially dehydrated, seeded with 1×10^5^ B16-F10 mouse melanoma cells, and incubated at 37 °C for 24 h in normoxia (Figure 3e and 3g) or hypoxia (Figure 3f and 3g). In normoxia (Figure 3e), B16-F10 cell-laden cryogels containing up to 0.5% CaO_2_ displayed high viability (∼97%), similar to their cryogel analogues, indicating their cytocompatibility. Cryogels containing 1% CaO_2_ showed a slight decrease in viability (85 ± 2%). However, regardless of the O_2_-cryogel composition, cells homogeneously attached and spread throughout the constructs (Figure 3g). When incubated in hypoxia, similar results were obtained in terms of cell attachment and spreading (Figure 3g), but the viability of B16-F10 cells remained high (95 ± 2%) throughout the assessed CaO_2_ concentration range (Figure 3f). These results suggest that the O_2_-cryogels developed in this study are cytocompatible, promote cell adhesion, and support cell viability under both normoxic and hypoxic conditions.

To further investigate the biocompatibility of O_2_-cryogels, APC-free and APC-functionalized O_2_-cryogels containing 1% CaO_2_ were injected into the subcutaneous space of the dorsal flanks of C57BL/6J mice. Cryogels functionalized with APC were used as controls. Seven days post-injection, the cryogels and O_2_-cryogels were excised with the surrounding tissue and stained using H&E and MT for histological analysis (Figure 3h). All explanted biomaterials retained their initial size. Only a few neutrophils were observed in the APC-functionalized cryogels and O_2_-cryogels, suggesting minimal inflammation. Furthermore, the junction between the connective tissue stroma and these cryogels consisted mainly of fibrin, suggesting that the biomaterials were well integrated within the native tissue. In contrast, APC-free O_2_-cryogels showed a high number of neutrophils in the connective tissue stroma and within the implant, indicating an inflammatory reaction. Collectively, these results suggest that O_2_-cryogels are biocompatible and that the enzyme CAT contributes to preventing a host inflammatory response.

### Reversing local cellular tumor hypoxia and associated immunosuppression

O_2_-cryogels were next analyzed for their capacity to reverse hypoxia at the cellular level. Cellular hypoxia of B16-F10 in cryogels at various amounts of CaO_2_ was assessed in the presence of Hypoxyprobe™-1 (Figure 4a,b). B16-F10 cells (1×10^5^) were seeded in cryogels or O_2_-cryogels and incubated at 37 °C for 24 h in hypoxia. After incubation, 96 ± 1.2% of the cells within cryogels became hypoxic. Higher CaO_2_ concentrations within O_2_-cryogels were associated with lower percentages of hypoxic cells, between 84.5% (with 0.1% CaO_2_) and 10.6% (with 1% CaO_2_). Additionally, no significant difference was observed between cells within O_2_-cryogels containing 1% CaO_2_ cultured in hypoxia and cells within cryogels cultured in normoxia, suggesting effective reversal of cellular hypoxia.

In addition to analyzing cellular hypoxia, the concentration of extracellular adenosine − downstream of hypoxia − was measured within the supernatant of cell-laden cryogels and O_2_-cryogels after 24 h of incubation (Figure 4c). As expected, cells cultured within cryogels and subjected to hypoxia showed increased levels of extracellular adenosine (0.85 ± 0.05 μM) compared to those cultured in normoxic conditions (0.68 ± 0.08 μM). However, in hypoxia, O_2_-cryogels were associated with significant decreased levels of extracellular adenosine, with values similar to those in normoxic conditions obtained with 0.5% CaO_2_ (0.67 ± 0.05 μM) and 1% CaO_2_ (0.65 ± 0.06 μM).

The effect of local oxygenation on hypoxic B16-F10 cell phenotype was also investigated. A sudden change in oxygen concentration or cell toxicity can impact cell phenotype. In this context, only O_2_-cryogels containing 0.5% CaO_2_, a formulation allowing sustained release of oxygen over 30 h, were tested and compared to their CaO_2_-free cryogel analogues. B16-F10 cells were seeded within these cryogels and cultured under normoxic or hypoxic conditions for 24 h. Total mRNA was then extracted from the cells, and the expression profile of the hypoxia-inducible genes *HIF1α, Wnt11, CD73, VEGFα, CD44, CD73*, and *CD133* was analyzed by RT-qPCR (Figure 4d). As expected, when seeded in cryogels under hypoxic conditions, B16-F10 cells demonstrated increased expression of *HIF1α* (+45%), *Wnt11* (+123%), *CD73* (+75%), *VEGFα* (+125%), *CD44* (+140%), and *CD133* (+246%) compared to the expression levels under normoxic conditions. In contrast, no changes in gene expression were observed in cells seeded in O_2_-cryogels containing 0.5% CaO_2_ under hypoxic conditions. Additionally, in comparison to cells cultured in hypoxia with cryogels, *CD44* and *CD133* gene expression levels decreased in B16-F10 cells cultured in O_2_-cryogels (by 65% and 48%, respectively). Collectively, these results suggest that O_2_-cryogels can reverse cellular hypoxia, thereby preventing the accumulation of immunosuppressive adenosine and the induction of a hypoxic and aggressive phenotype in cancer cells.

### Restored cytotoxicity of hypoxic OT-1 T cells in an antigen-specific B16-OVA model

Several studies have established that the hypoxic TME dramatically impairs the ability of tumor-reactive T cells to eliminate tumor cells.(*16, 17, 54*) Thus, we investigated the effect of O_2_-cryogels on the tumoricidal capacity of spleen-derived OVA-specific OT-1 T cells to eliminate B16-OVA cells in normoxic and hypoxic culture (Figure 5a). Cytotoxic T-cell function was first assessed in real time by time-lapse microscopy (Figure 5b and Supplementary Videos 4,5). B16-OVA cells were cultured for 3 h in 24-well plates in normoxia to allow adhesion and subsequently cocultured with OT-1 T cells for 4 h in hypoxia in the presence of cryogels or O_2_-cryogels containing 0.5% CaO_2_. As expected, under hypoxic conditions, antigen-specific OT-1 T cells did not induce tumor cell death when exposed to cryogels (Figure 5b and Supplementary Video 4). However, when incubated with O_2_-cryogels, the defective cytotoxic function of hypoxic OT-1 T cells was rescued, leading to the rapid killing (< 4 h) of B16-OVA tumor cells (Figure 5b and Supplementary Video 5).

**Figure 5.**
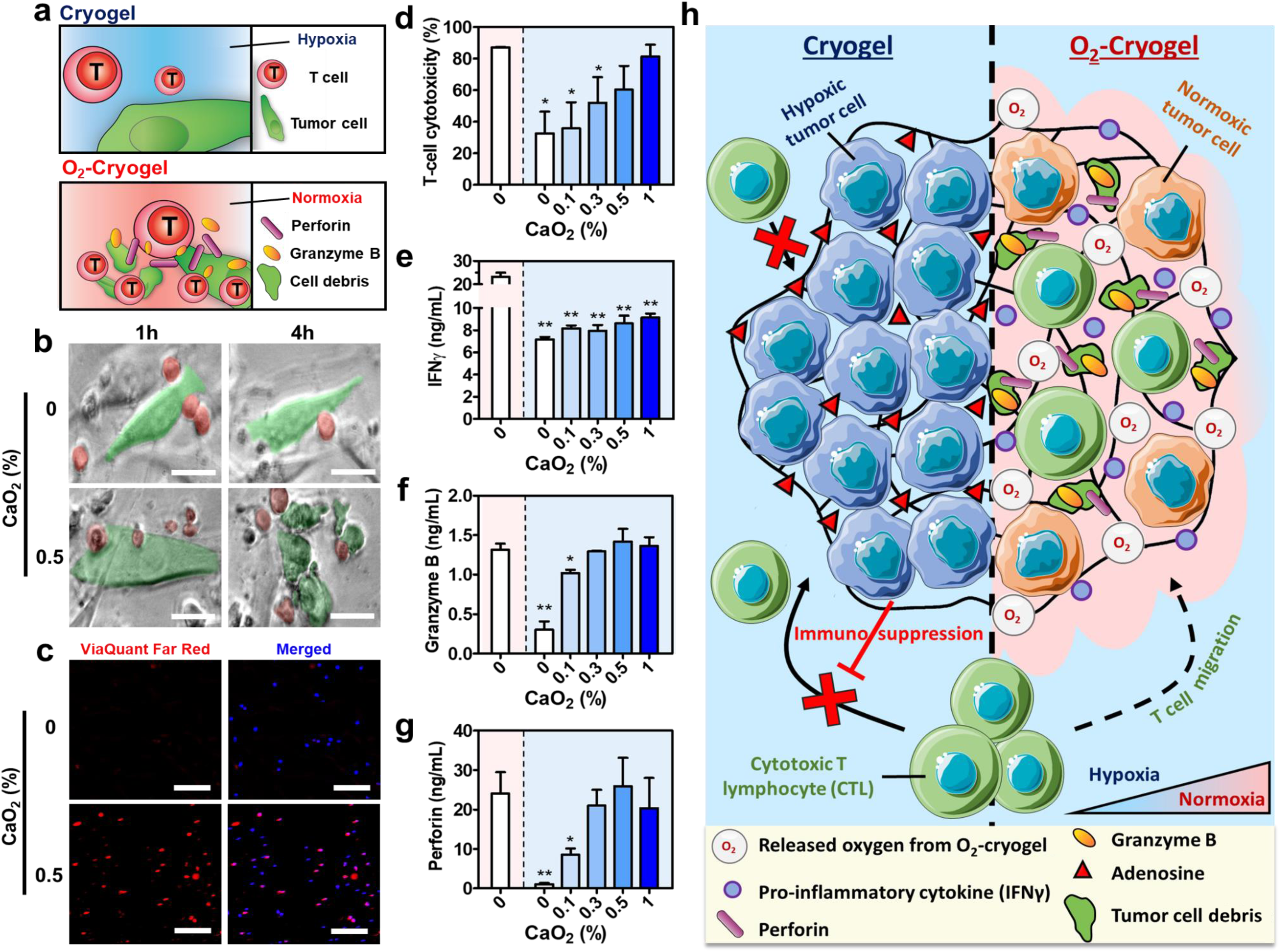
O_2_-cryogels restore T cell-mediated cytotoxicity against hypoxic tumor cells. **a**) Restored T cell activity in O_2_-cryogels. **b**) Time-lapse images showing OT-1 T cells attacking B16-OVA cells in cryogels and O_2_-cryogels containing 0.5% CaO_2_ after incubation for 1 h (top) and 4 h (bottom) under hypoxic conditions (1% O_2_). B16-OVA cells and OT-1 T cells are pseudocolored in green and red, respectively. **c**) Confocal images showing cell viability of B16-OVA in O_2_-cryogels containing 0% or 0.5% CaO_2_ after 24 h of co-culture with OT-1 T cells under hypoxic conditions (1% O_2_). Red = dead cells stained with ViaQuant Far Red, blue = nuclei stained with DAPI. The data are representative of n = 5 samples per condition. d-g) B16-OVA cells were cocultured with OT-1 T cells for 24 h under normoxic (20% O_2_, red background) or hypoxic (1% O_2_, blue background) conditions. **d**) Quantification of T cell-mediated cytotoxicity. **e–g**) Secretion of IFNγ (e), granzyme B (f), and perforin (g) from OT-1 T cells. **h**) Schematic depicting an O_2_-cryogel (*i*) inducing local oxygenation in an oxygen-deprived solid tumor, (*ii*) reversing hypoxia-driven immunosuppression, (*iii*) restoring the tumoricidal function of cytotoxic T cells fatal to tumor cells. Values represent the mean ± SEM (n = 4). Data were analyzed using ANOVA and Dunnett’s post-hoc test (compared to cryogels), ^*^P < 0.5, ^**^P < 0.01. Scale bars = 20 µm (b) and 100 µm (c).

To substantiate these observations, we compared the effect of the CaO_2_ concentration on T-cell cytotoxicity in hypoxic and normoxic conditions (Figure 5c–d and Supplementary Figures 5,6a). To this end, B16-OVA cells were first cultured in cryogels containing different CaO_2_ concentrations for 3 h in normoxia and then coincubated with OT-1 splenocytes for 4 and 24 h in normoxia or hypoxia. Under normoxic conditions, cytotoxic OT-1 T cells infiltrated O_2_-cryogels and killed B16-OVA cells (85 ± 5% cytotoxicity) within 4 h (Supplementary Figure 6a), regardless of the CaO_2_ concentration. In hypoxic conditions, the cytotoxic activity of OT-1 T cells within cryogels decreased substantially (4 h = 3.55 ± 1%; 24 h = 34.38 ± 11% cytotoxicity), confirming that low oxygen tension inhibits the cytotoxicity of T cells in this model (Figure 5c,d and Supplementary Figure 6a); this is consistent with the time-lapse microscopy results. After 4 h of coincubation in hypoxia, O_2_-cryogels restored the OT-1 T-cell cytotoxicity by up to 50% in a CaO_2_ concentration-dependent manner (Supplementary Figure 6a). Moreover, O_2_-cryogels containing 0.5% and 1% CaO_2_ particles fully restored the cytotoxicity of OT-1 T cells after 24 h in hypoxia to similar levels to those in cells cultured in normoxia (Figure 5c,d).

**Figure 6:**
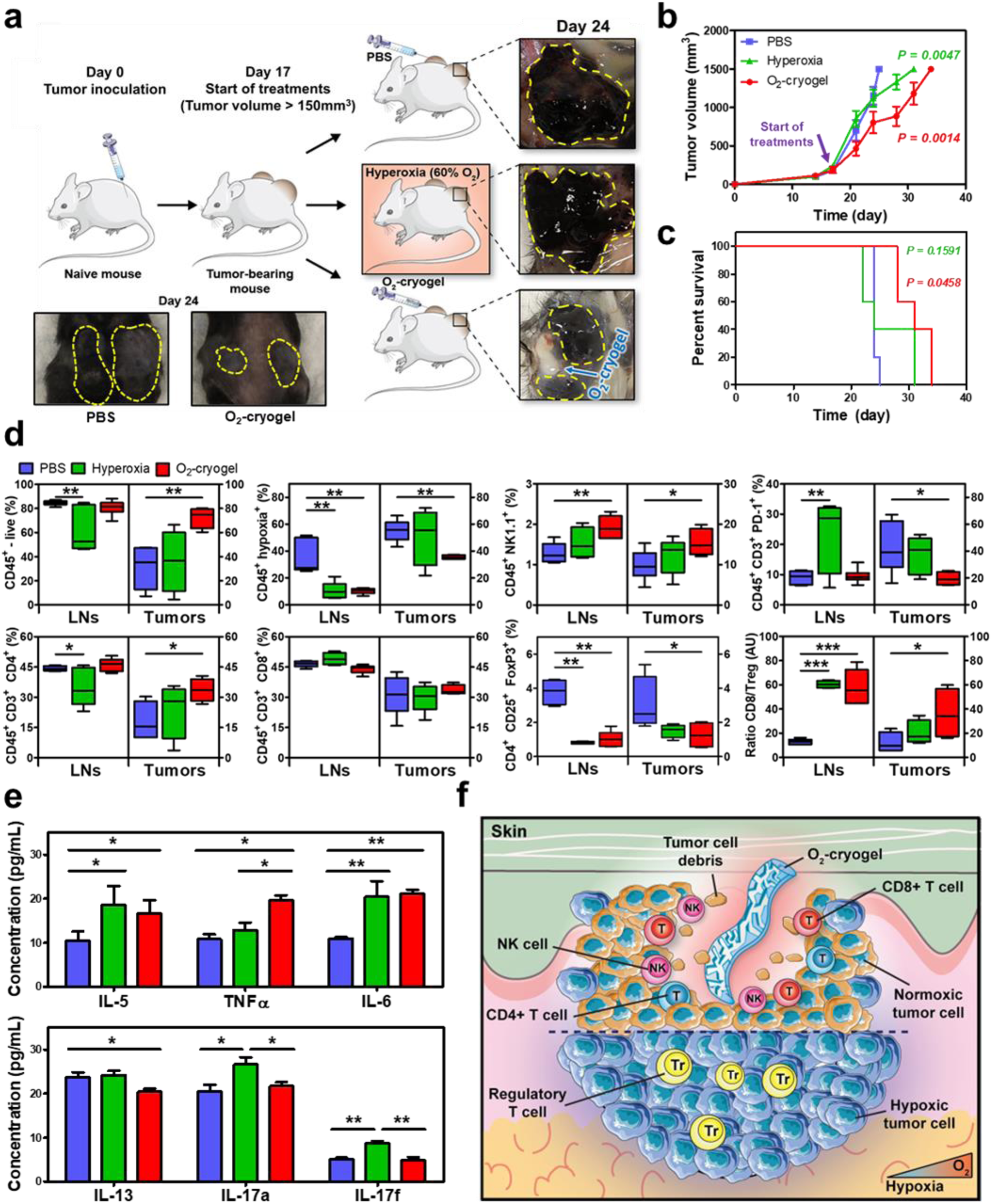
Antitumor immune response and survival in advanced-stage tumor-bearing mice treated with O_2_-cryogel therapy as co-adjuvant. **a**) Mice were inoculated at day 0 with 2×10^5^ B16-F10 cells in both flanks. On day 17, when the tumor volume reached > 150 mm^3^, tumor-bearing mice were injected intratumorally with PBS or O_2_-cryogels twice a week until the end of the study (n = 10/group). Another group of tumor-bearing mice (n = 10) was placed into a hyperoxic chamber at 60% O_2_. On day 24, half of the mice were sacrificed to perform mechanistic studies (n = 5/group, mouse back and tumor photographs), and the other half (n = 5/group) continued treatment. **b**) Melanoma tumor growth curve. **c**) Kaplan-Meier survival curves based on a 1,500 mm^3^ tumor volume as the endpoint criteria. **d**) Flow cytometry quantification of the number of immune cells (CD45^+^), hypoxic immune cells (CD45^+^ Hypoxia^+^), natural killer cells (CD45^+^ NK1.1^+^), PD-1 positive T cells (CD45^+^ CD3^+^ PD-1^+^), T helper cells (CD45^+^ CD3^+^ CD4^+^), cytotoxic T cells (CD45^+^ CD3^+^ CD8^+^), and regulatory T cells (CD45^+^ CD3^+^ CD4^+^ CD25^+^ FoxP3^+^), as well as the ratio of CD8^+^/regulatory T cells in the tumor-draining inguinal lymph nodes (LNs) and tumors at day 24 (i.e., day 7 post-treatment). **e**) Quantification of anti-tumor (IL-5, IL-6, TNF-α) and pro-tumor (IL-13, IL-17a, IL-17f) serum cytokines at day 7 post-treatment. **f**) Schematic illustrating the O_2_-cryogel therapeutic approach in solid tumors. In hypoxic tumors (bottom), the immunosuppressive environment prevents cytotoxic T cell and NK cell recruitment and function. O_2_-cryogels (top) reverse local hypoxia, leading to increased NK, T helper, and cytotoxic T cell infiltration and function. Values represent the mean ± SEM (n = 5 mice per group). Data were analyzed using a two-tailed Pearson test (b), a two-tailed Mantel-Cox text (c), and an ANOVA with a Dunnett’s post-hoc test (compared to PBS), ^*^P < 0.05, ^**^P < 0.01, ^***^P < 0.001.

This finding was further confirmed by evaluating the concentrations of IFNγ, perforin, and granzyme B secreted by functional OT-1 T cells after 4 h (Supplementary Figure 5b–d) and 24 h (Figure 5e–g). IFNγ is a key mediator of cytotoxic T cell motility and cytotoxicity,^(*55*)^ and perforin and granzyme B are the main molecules involved in inducing apoptosis and mediating cytotoxicity of the B16-OVA target cells.(*56*) The IFNγ, perforin, and granzyme B concentrations were dramatically lower in the supernatant of cell-laden cryogels in hypoxia (60% to 95% inhibition) than in their normoxic counterparts, in agreement with the cytotoxicity assessments. While O_2_-cryogels containing 0.1% CaO_2_ had a minimal impact on hypoxic OT-1 T cells within the first 4 h, they were able to partially relieve the hypoxic stress after 24 h, resulting in a slight increase in perforin and granzyme B production. Strikingly, cryogels containing higher amounts of CaO_2_ completely restored the ability of OT-1 T cells to secrete perforin and granzyme B within 24 h. Overall, these results indicate that O_2_-cryogels can locally reverse hypoxia, overcome hypoxia-driven T-cell inhibition, and restore the killing ability of OT-1 T cells via the secretion of proinflammatory cytokines and cytotoxic proteins fatal to B16-OVA cells (Figure 5h).

### Boosting antitumor immune responses in an aggressive and advanced murine melanoma model

After demonstrating that O_2_-cryogels reversed hypoxia-induced immunosuppression in vitro, we next tested their capacity to promote antitumor immunity as a co-adjuvant in a highly aggressive and advanced-stage murine melanoma model (Figure 6a). Advanced tumors were first established via subcutaneous inoculation of 2×10^5^ B16-F10 tumor cells on both flanks, and treatments were administered once advanced-stage tumors reached > 150 mm^3^ in volume (on day 17). PBS or O_2_-cryogel were injected in each tumor (2 tumors/mouse), and as a control group, mice were placed in a hyperoxic chamber at 60% oxygen to promote immune-mediated tumor rejection.(*17*) At day 24, photographs of subcutaneous tumors and excised tumors showed that the size of O_2_-cryogel-treated tumors was significantly decreased compared to tumors in mice treated with PBS (sham) or respiratory hyperoxia. Strikingly, we observed local eradication of tumor tissue around O_2_-cryogels after excision. Tumor growth for mice treated with hyperoxia was slightly reduced compared to PBS, whereas tumor sizes were markedly smaller following intratumoral injections of O_2_-cryogels (Figure 6b). As a result, O_2_-cryogel treatment extended mouse survival up to 35 days, with a median survival time of 31 days as opposed to 23 days for PBS or respiratory hyperoxia (Figure 6c). Overall, in comparison to PBS, the median lifespan of treated mice with O_2_-cryogels was extended by nearly 35%.

To understand the mechanisms underlying the improved antitumor immune responses in O_2_-cryogel-treated mice, immune cell types and fractions were quantified in tumors, tumor-draining LNs, and spleens at day 24 (Figure 6d and Supplementary Figure 7). While the fraction of viable immune cells was only 31 ± 8 % in PBS-treated tumors, O_2_-cryogels greatly increased the viable immune cell fraction to 72 ± 4 % (Figure 6d). Surprisingly, immune cell viability in the inguinal draining LNs was reduced in hyperoxia-treated mice. Furthermore, O_2_-cryogels reduced the fraction of hypoxic immune cells in tumors, LNs, and spleens by 50% relative to PBS-injected tumors (Figure 6d, Supplementary Figure 7). The capacity of O_2_-cryogels to promote antitumor immunity is further highlighted by the increased fractions of tumor-infiltrating NK cells from 9% to 15% and CD4^+^ T cells from 18% to 33%. In tumor-draining LNs, the number of NK cells was also increased (1.2% to 1.9%) with O_2_-cryogels, whereas the number of CD4^+^ T cells did not. In contrast, hyperoxia significantly reduced CD4^+^ T-cell numbers in the LNs. Surprisingly, the fractions of CD8^+^ T cells in tumors, LNs, and spleens were not impacted by either O_2_-cryogel or respiratory hyperoxia. O_2_-cryogels were also shown to counteract immunosuppression in tumors and LNs by considerably reducing the expression of co-inhibitory receptor PD-1 on intratumoral T cells from 42% to 19%. Although O_2_-cryogel injection did not alter PD-1 expression on T cells in the LNs and spleens, we observed an increase in the fraction of PD-1^+^ T cells in the LNs and spleens from hyperoxia-treated mice. Unlike with PBS, mice treated with hyperoxia and O_2_-cryogels exhibited a substantial reduction in the fractions of CD4^+^CD25^+^FoxP3^+^ regulatory T cells in tumors and LNs, leading to higher ratios of cytotoxic CD8^+^ T cell to regulatory T cell by three-to four-fold, respectively.

To further characterize the oxygen-mediated immune response, serum cytokine levels were assessed at day 24 (Figure 6e). Compared to PBS, hyperoxia and O_2_-cryogel treatment resulted in a two-fold increase in the concentrations of pro-inflammatory cytokines IL-5 and IL-6. There was also a marked increase in serum TNFα from 10.8 ± 1 to 19.6 ± 1.1 pg/mL with O_2_-cryogels. Levels of several anti-inflammatory or tumor-promoting cytokines were also altered after treatments. Interestingly, intratumoral O_2_-cryogel injection reduced serum levels of tumor-promoting cytokine IL-13 from 23.7 ± 1.1 to 20.4 ± 0.6 pg/mL. Moreover, IL-17 concentrations were increased in hyperoxia-treated mice. Serum levels of IFNγ, IL-2, IL-4, IL-9, and IL-22 were unaltered after hyperoxia or O_2_-cryogel treatment (Supplementary Figure 8). Taken together, these data support a model in which O_2_-cryogels act as a co-adjuvant to reverse intratumor hypoxia-mediated immunosuppression in vivo (Figure 6f). This strategy reduces the number of regulatory T cells and expression of immune checkpoints. Furthermore, local oxygenation increases infiltration of cancer-fighting immune cells, including T cells and NK cells, leading to local elimination of tumor tissues.

## Discussion

Hypoxia plays a key role in both the development and maintenance of tumor immune privilege. To overcome hypoxia-mediated tumor resistance, approaches targeting hypoxia, such as systemic hyperbaric oxygen therapy, and hypoxia-inducible signaling pathways such as HIF inhibitors and A2AR antagonists, have shown encouraging results in preclinical and clinical studies.(*30-33*) These approaches have been reported to boost tumor-specific cytotoxicity of T cells and NK cells.(*17*) However, to date, the modest efficacy in melanoma coupled with potential treatment-related toxicities have prevented their implementation as a standard of care.(*57*) Therefore, it is important to have an alternative, non-systemic anti-hypoxic strategy to reduce tumor hypoxia and enhance cancer immunotherapies. To this end, we developed a new and promising approach by engineering O_2_-cryogels to target oxygen delivery in oxygen-deprived environments.

We created a new class of sponge-like and syringe-injectable O_2_-cryogels that can produce oxygen in a controlled and sustained fashion, enabling reversal of local tumor hypoxia and restoration of T cell-mediated cytotoxicity. These O_2_-cryogels were fabricated by physically caging solid CaO_2_ particles within the cross-linked polymer network of cryogels. CaO_2_ particles were homogenously distributed throughout the cryogel structure, providing predictable and spatiotemporally controllable release of oxygen. To prevent H_2_O_2_-related toxicity, chemically modified CAT (APC) was covalently incorporated to the cryogel backbone.

Oxygen diffusion is often geometrically restricted in conventional (i.e., mesoporous) hydrogels,(*58*) altering both the cellular and local tissue oxygenation.(*40, 43*) However, from O_2_-cryogels, oxygen is freely released to the surrounding environment, reaching up to 800 μM after 2 h, and in a sustained fashion for 2 days. O_2_-cryogel-mediated oxygenation reversed cellular hypoxia for at least 24 h, both through the entire construct and to distant locations. The highly interconnected macroporous channels of O_2_-cryogels facilitate oxygen transport into the surrounding environment, resulting in increased oxygen levels. In this scenario, oxygen release relies on the diffusion of free water through the polymer network to hydrolyze CaO_2_ particles. However, in body tissues, bound water may result in slower and more sustained release of oxygen. Therefore, oxygen release kinetics could be further optimized for in vivo applications.(*59-61*) For instance, encapsulation of perfluorocarbon or H_2_O_2_/polyvinylpyrrolidone (PVP)(*62, 63*) particles in O_2_-cryogels could facilitate oxygen production in body tissues. Additionally, other peroxides (e.g., magnesium peroxide, zinc peroxide), oxides (e.g., manganese dioxide and zinc oxide), and percarbonates (e.g., sodium percarbonate) could be used as alternative oxygen-generating agents.^(*43*)^

O_2_-cryogels were conjugated with CAT to decompose H_2_O_2_ into additional oxygen. This feature resulted in a safe biomaterial when subcutaneously injected in mice, and cells cultured within these constructs remained viable under both normoxic and hypoxic conditions. However, cell viability was slightly altered when B16-F10 cells were cultured in O_2_-cryogels containing 1% CaO_2_. This could be attributed to a partial loss of enzymatic activity during the chemical modification of CAT and its successive cryogenic treatments. Furthermore, once anchored within the polymer walls, the steric hindrance could also alter CAT activity in depleting H_2_O_2_. To protect CAT, several techniques could be investigated. For instance, optimizing the chemical modification of CAT, introducing different PEG spacers to reduce steric hindrance, and/or incorporating lyoprotectants (e.g., trehalose) to prevent freeze-drying induced protein denaturation should help preserve CAT to deplete H_2_O_2_ more effectively.(*64*) Alternatively, conjugating CAT to cryogels post-fabrication could also be used as a means to retain its enzymatic activity.

Oxygen-generating biomaterials have the potential to restore the function of impaired pre-existing T cells when subjected to hypoxia. In this work, we showed that O_2_-cryogel-induced oxygenation can reinstate T cell-mediated antitumor activity in hypoxia to levels comparable to those observed in normoxia. Although there was only a slight increase in IFNγ secretion, hypoxia reduction substantially enhanced T cell-mediated secretion of cytotoxic proteins such as perforin and granzyme B. Strikingly, the tumoricidal function of T cells was restored after only 4 h, and as little as 0.1% CaO_2_ was sufficient to boost cytotoxic protein secretion after 24 h. Overcoming cellular hypoxia most likely reinforced the killing ability of OT-1 T cells against tumor cells. Mechanistically, O_2_-cryogels appeared to prevent stabilization of hypoxia-dependent HIF1α, a major orchestrator of tumor cell adaptation to hypoxia and aggressiveness. O_2_-cryogels also induce the downregulation of VEGFα, Wnt11, CD44, and CD133, which have been associated with angiogenesis as well as tumor cell proliferation, migration, and invasion in solid tumors.(*65-69*) Additionally, O_2_-cryogels inhibited the expression of adenosine-producing ecto-enzyme CD73. This was further confirmed with reduced concentrations of extracellular adenosine, a key immunosuppressive metabolite that may allow “cold” tumors to escape immune surveillance.(*25-27, 70*) Collectively, our data suggest that O_2_-cryogels contribute to oxygen-mediated restoration of T-cell effector function within a hypoxic TME. This is in agreement with previous studies where Sitkovsky et al. showed that oxygenation increases the density of peptide-major histocompatibility complex (MHC) class I molecules at the surface of tumor cells, enhancing their recognition and elimination by tumor-specific T cells.(*71*)

O_2_-cryogels developed in this study can withstand the injection process without releasing CaO_2_ particles. Furthermore, these gels can overcome shear stress during injection while retaining their oxygen-generating feature. This characteristic is a significant advantage over previously reported oxygen-generating preformed hydrogels that require invasive surgical implantations.(*41, 42, 46*) We believe that, similarly to systemic oxygenation,(*72*) local delivery of oxygen from O_2_-cryogels can be used as co-adjuvant therapy in the treatment of malignancies. However, unlike systemic oxygenation, local oxygen delivery from O_2_-cryogels has the potential to target any solid tumors and prevent potential side effects (e.g., pulmonary edema).(*73*) In this work, we showed that respiratory hyperoxia, our positive control, and O_2_-cryogel co-adjuvant therapies can both slow down tumor growth and increase mice survival in a highly aggressive and advanced-stage murine melanoma model. Both strategies reduced the fractions of hypoxic immune cells in tumors, allowing these cells to overcome immunosuppression(*74*), as indicated with the decrease of tumor-infiltrating regulatory T cells. However, in our study, respiratory hyperoxia was associated with reduced immune cell viability in the tumor-draining inguinal LNs, and PD-1 expression on T cells was increased, suggesting that long-term systemic oxygenation and/or poor oxygenation of subcutaneous tumors could impair immune cell function. In addition, this treatment led to high serum levels of IL-17 and IL-13, two cytokines promoting cancer progression,(*75, 76*) which is in agreement with the moderate effect seen with respiratory hyperoxia on curbing tumor growth. In contrast, O_2_-cryogel therapy increased CD4^+^ T cell number and the CD8^+^/regulatory T cell ratio in tumors, suggesting a significant shift in the immune balance from immunosuppression to active antitumor immunity. This is further demonstrated by elevated serum level of TNF-α, indicating that local oxygen delivery mediated a strong pro-inflammatory immune response. Interestingly, O_2_-cryogels also enhanced immune cell survival within the tumors and promoted infiltration of NK cells, important contributors to antibody-dependent cell-mediated toxicity and cell-mediated immune responses against cancer.(*77*) These observations might explain why O_2_-cryogel-mediated co-adjuvanted oxygen therapy induced tumor regression locally, delayed overall tumor growth, and prolonged survival. Collectively, our data suggest that O_2_-cryogel therapy outperformed respiratory hyperoxia, used here as our gold standard. Furthermore, local oxygen delivery may provide a therapeutic strategy to reduce hypoxic regions within solid tumors across various cancers and drive an influx of cancer-fighting effector cells(*17*) to induce tumor regression or rejection.(*16, 22, 46*)

Recently, injectable cryogel vaccines have shown to be effective as a prophylactic treatment against melanoma.(*78, 79*) These vaccines elicit strong and long-lasting antitumor immune responses. Yet, their efficacy in a therapeutic setting is limited and requires further optimization. In this context, O_2_-cryogels could potentially be used as co-adjuvant to reinforce DC activation and ultimately induce downstream stimulation of cytotoxic T cells. Additionally, further in vivo testing is required to fully assess the clinical potential of our approach and investigate whether O_2_-cryogels can rescue other oxygen-deprived antitumor immune cells, such as macrophages and dendritic cells.

Moving forward, innovative biomaterials such as O_2-_cryogels hold great potential to advance the future of cancer immunotherapy.(*49, 50, 78-82*) Considering the recent clinical approval of several immune checkpoint inhibitors for cancer treatment(*83*) and the recent advances in targeting the hypoxia-adenosinergic signaling pathway,(*84*) the combination of O_2_-cryogels with immune checkpoint blockade (e.g., CTLA-4 or PD-1)(*17*) should be investigated for a synergistic effect. This is further supported with recent studies suggesting hypoxia as a potential obstacle on immune checkpoint inhibitors to treat cancer.(*17, 85*) Additionally, O_2_-cryogel-mediated hypoxia reduction could be combined with other strategies under development to tackle hypoxia-driven immunosuppression such as A2AR antagonists,(*31, 86*) HIF inhibitors,(*30, 33*) and hyperoxia(*16, 34*) to cooperatively bypass hypoxia-driven T-cell immunosuppression and restore tumor-reactive T-cell function.

## Methods

### Materials

Sodium salt of HA, phosphate-buffered saline (PBS), dimethylformamide (DMF), glycidyl methacrylate (GM), triethylamine (TEA), adipic acid dihydrazide (AAD), 1-ethyl-3-(3-dimethylaminopropyl)-carbodiimide (EDC), CAT, calcium peroxide (CaO_2_) particles, tetramethylethylenediamine (TEMED), ammonium persulfate (APS), Alizarin Red S, fluorimetric hydrogen peroxide assay kit, fetal bovine serum (FBS), paraformaldehyde (PFA), Triton X-100, and 4′,6-diamidino-2-phenylindole (DAPI) were purchased from Sigma-Aldrich (St. Louis, MO, USA). NHS-rhodamine, 5/6-carboxy-tetramethyl-rhodamine succinimidyl ester (NHS-rhodamine), penicillin, streptomycin, gentamycin, rabbit anti-mouse IgG (H+L) Alexa Fluor 488 secondary antibody, PureLinkTM RNA Mini Kit, High Capacity cDNA Reverse Transcription Kit gene expression assays were purchased from Thermo Fisher Scientific (Waltham, MA, USA). IFNγ, TNFα, and granzyme B DuoSet ELISA kits were purchased from R&D Systems (Minneapolis, MN, USA). Carboxyfluorescein succinimidyl ester (CFSE), CD45 (PE/Cy7, clone 104), NK1.1 (APC, clone PK136), PD-1 (BV421, clone 29F.1A12), CD3 (BV605, clone 17A2), CD4 (PerCP/Cy5.5, clone GK1.5), CD8 (Alexa Fluor 700, clone 53-6.7), CD25 (APC/Cy7, clone 3C7) and FoxP3 (PE, clone MF-14) antibodies, and LEGENDplex Mouse Th Cytokine Panel were purchased from Biolegend (San Diego, CA, USA). OVA 257-264 (SIINFEKL peptides) was purchased from InvivoGen (San Diego, CA, USA). Amine-terminated GGGGRGDSP (G_4_RGDSP) peptide was purchased from Peptide 2.0 (Chantilly, VA, USA). Acrylate-PEG-N-hydroxysuccinimide (3.5 kDa) was purchased from JenKem Technology (Plano, TX, USA). ViaQuant™ Fixable Far-Red Dead Cell Staining Kit was purchased from GeneCopoeia (Rockville, MD, USA). Alexa Fluor 488-phalloidin was purchased from Cell Signaling Technology (Danvers, MA, USA). The Hypoxyprobe Plus kit was purchased from Hypoxyprobe Inc. (Burlington, MA, USA). A fluorometric adenosine assay kit was purchased from Biovision (San Francisco, California, USA). The Perforin ELISA kit was purchased from G-Biosciences (St. Louis, MO, USA). Melanoma cells (B16-F10, CRL-6475) were purchased from ATCC (Rockville, MD, USA).

### Chemical modification of hyaluronic acid

HA was conjugated with GM as previously described.(*87*) Briefly, HA salt (5 g) was dissolved in PBS (1 L, pH 7.4) and mixed with DMF (335 mL), GM (62 mL), and TEA (46 mL). The reaction was allowed to proceed for ten days at room temperature (RT), then the mixture was precipitated in a large excess of acetone, filtered using grade 4 Whatman paper, and dried overnight at RT in a vacuum oven. The resulting product, hyaluronic acid glycidyl methacrylate (HAGM), was characterized by ^1^H nuclear magnetic resonance (NMR).

Rhodamine-labeled HAGM (R-HAGM) was prepared by coupling NHS-Rhodamine with amine-terminated HAGM (NH_2_-HAGM) as previously described.(*87*) To synthesize NH_2_-HAGM, HAGM (1 g) was dissolved in 0.1 M 2-(N-morpholino) ethanesulfonic acid (MES) (100 mL) at pH 5.5. Next, AAD (4 g) and EDC (90 mg) were added to the solution. The reaction mixture was stirred at RT for 4 h to allow the amination of HAGM to proceed. NH_2_-HAGM was then precipitated in a large excess of acetone, filtered, and dried in a vacuum oven overnight at RT. NH_2_-HAGM (1 g) was dissolved in 0.1 M NaHCO_3_ buffer solution (10 mL) at pH 8.5 and mixed with NHS-Rhodamine (10 mg). The reaction was allowed to proceed for 16 h at RT, and R-HAGM was precipitated in a large excess of acetone, filtered, and then dried overnight at RT in a vacuum oven.

### Chemical modification of G_4_RGDSP peptide and catalase

Acrylate-PEG-G_4_RGDSP (APR) was synthesized by coupling the amine-terminated G_4_RGDSP peptide to acrylate-PEG-N-hydroxysuccinimide (molar ratio, 1:1). Briefly, acrylate-PEG-N-hydroxysuccinimide (100 mg) and G_4_RGDSP (22.3 mg) were mixed in 1 mL 0.1 M NaHCO_3_ buffer solution at pH 8.5, allowed to react for 4 h at RT, and freeze dried overnight. Similarly, acrylate-PEG-CAT (APC) was synthesized by coupling the CAT enzyme to acrylate-PEG-N-hydroxysuccinimide comonomers (molar ratio, 1:3).

### Fabrication of O_2_-cryogels

CaO_2_-containing cryogels (i.e., O_2_-cryogels) were fabricated by redox-induced free radical cryopolymerization in dH_2_O at −20 °C. Briefly, HAGM (40 mg, 4% wt/vol) was mixed with APR (8 mg, 0.8% wt/vol), APC (10 mg, 1% wt/vol), and various concentrations of CaO_2_ (0**–**1% wt/vol) in dH_2_O (1 mL). The mixture was precooled at 4 °C, and then TEMED (0.42% wt/vol) and APS (0.84% wt/vol) were added to the solution. The solution was poured into Teflon® molds, transferred to a freezer preset at −20 °C, and cryopolymerized for 16 h. Finally, O_2_-cryogels were brought to RT to remove ice crystals and washed with dH_2_O. For microscopy imaging, a fraction of HAGM (0.3% wt/vol) in each formulation was substituted with R-HAGM to fluorescently stain the polymer network of cryogels.

### Characterization of CaO_2_ encapsulation and distribution, and O_2_-cryogel microstructural features

The encapsulation efficiency of CaO_2_ within the polymer walls of O_2_-cryogels was determined by Alizarin Red S staining, which stains red on reaction with inorganic calcium. Specifically, a 2% wt/vol Alizarin Red S solution was freshly prepared in dH_2_O and adjusted to pH 4.2. O_2_-cryogels were incubated with Alizarin Red S for 20 min, rinsed with dH_2_O several times until the washing solution was clear, and photographed with a Canon camera or imaged using a Leica TCS SP5 X WLL confocal microscope (Buffalo Grove, IL, USA). The CaO_2_ distribution in the polymer walls as well as the microstructural features of O_2_-cryogels were characterized by scanning electron microscopy and energy-dispersive X-ray analysis (EDX). First, O_2_-cryogels were freeze dried for 24 h, mounted on a sample holder using carbon tape, and sputter coated with platinum/palladium up to a thickness of 5 nm. The resulting samples were imaged by secondary electron detection on a Hitachi S-4800 scanning electron microscope at a voltage of 25 kV and a current of 10 µA. Pore sizes were determined using the Fiji(*88*), Analyze Particles, and Diameter J software plugins.

### Physical characterization of O_2_-cryogels

The swelling ratio was determined using a conventional gravimetric procedure as previously described.(*49*) Briefly, O_2_-cryogel cylinders (6-mm diameter, 6-mm height) were prepared, immersed in PBS for 24 h at 37 °C, and weighed (m_s_). Next, O_2_-cryogel cylinders were washed in dH_2_O, freeze dried, and weighed again (m_d_). Then, as shown in Equation 1, the equilibrium mass swelling ratio (Q_m_) was calculated by dividing the mass of fully swollen O_2_-cryogel (m_s_) by the mass of freeze-dried O_2_-cryogel (m_d_):

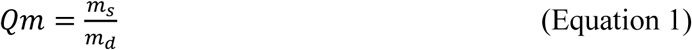

The degree of pore connectivity was evaluated using a water wicking technique. Specifically, fully hydrated O_2_-cryogel disks (6-mm diameter, 1-mm height) were first weighed on an analytical scale. Next, a Kimwipe® was used to wick away free water within interconnected pores, and gels were weighed a second time. The degree of pore connectivity was then calculated as the volume of water (%) wicked away from the gels.

Young’s modulus values were determined using an Instron 5944 universal testing system (Instron, Norwood, MA, USA). Cylindrical cryogels (6-mm diameter, 8-mm height) were dynamically deformed at a constant rate between two parallel plates for 10 cycles (strain rate of 10% per min) under constant hydration in PBS (pH 7.4). Compressive strain (mm) and load (N) values were then quantified at the 8th cycle using Instron’s Bluehill 3 software. Young’s modulus values were determined by measuring the tangent of the slope of the linear region on the loading stress-strain curve.

### Enzymatic activity of catalase and its derivatives (H_2_O_2_ depletion)

The enzymatic activities of CAT and APC were assessed using a fluorimetric hydrogen peroxide assay kit. Aqueous enzyme solutions were prepared at different concentrations (0.1–5 u/mL), mixed with 30 µM H_2_O_2_, and incubated at 37 °C for 30 min. Then, the fluorescence intensity (Ex/Em = 540/590 nm) was recorded using a Synergy HT plate reader (BioTek, Winooski, Vermont, USA) to quantify the residual H_2_O_2_ concentration in the solution. To evaluate potential H_2_O_2_ depletion from O_2_-cryogels, cryogel samples were incubated in dH_2_O at 37 °C for 15 min, transferred to 96-well plates, and assessed using the fluorimetric hydrogen peroxide assay kit, as described above.

### Oxygen release kinetics from O_2_-cryogels

The oxygen release kinetics of O_2_-cryogels were determined using needle-type optical oxygen microprobes (PyroScience GmbH, Aachen, Germany) under normoxic (20% O_2_) and hypoxic conditions (1% O_2_). Briefly, O_2_-cryogel cylinders (6-mm diameter, 8-mm height) were individually placed into 2 mL Eppendorf tubes containing Dulbecco’s modified Eagle’s medium (1.8 mL, DMEM) (Corning, Corning, NY, USA) and incubated at 37 °C under normoxic or hypoxic conditions using a Napco CO_2_ 1000 hypoxic incubator (Thermo Fisher Scientific). The needle-type probe was positioned in the center of the cylinders, and the dissolved oxygen concentration (µmol/L, 1 point every 300 seconds) was recorded for up to 2 days to monitor oxygen release. Additionally, to mimic syringe injection conditions, O_2_-cryogels were compressed by up to 90% of their volume prior to additional oxygen release measurements. Briefly, cryogel cylinders were allowed to swell in PBS until reaching equilibrium and were mechanically compressed between two parallel plates. The compressed cryogels were allowed to swell in DMEM until they returned to their initial shape and size, and oxygen release was monitored for up to 2 days at 37 °C under normoxic conditions.

### Cell seeding and culture in O_2_-cryogels

Melanoma cells (B16-F10) were cultured in complete DMEM (DMEM supplemented with 10% FBS, 100 µg/mL penicillin, and 100 µg/mL streptomycin). B16-F10 ovalbumin (OVA)-expressing cells (B16-OVA, kindly provided by Prof. M. Sitkovsky) were cultured in complete DMEM supplemented with 50 µg/mL gentamycin. Cells were incubated at 37 °C in either humidified 5% CO_2_/95% air (normoxia) or humidified 5% CO_2_/1% O_2_/94% N_2_ (hypoxia). Prior to cell seeding, square-shaped O_2_-cryogels (dimensions: 4 mm × 4 mm × 1 mm) were sanitized with 70% ethanol for 15 min, washed several times with sterile PBS, and mechanically compressed on sterile gauze to partially remove water from the pores under sterile conditions. Finally, a B16-F10 or B16-OVA cell suspension in complete DMEM (10^7^ cells/mL, 10 µL) was added dropwise to the top of each square-shaped cryogel and incubated for 1 h to allow cell adhesion. The cell-laden O_2_-cryogels were then supplemented with additional complete DMEM (2 mL) for the duration of the experiment. For experiments requiring a hypoxic environment, complete DMEM was preincubated (conditioned) for at least 1 h under hypoxic conditions.

### Cell viability assay

Cell viability within O_2_-cryogels was evaluated by a live/dead assay. After a 1-day incubation, cell-laden O_2_-cryogels were incubated for 15 min in the presence of a ViaQuant™ Fixable Far-Red Dead Cell Staining Kit according to the manufacturer’s instructions. Next, the cryogels were rinsed with PBS (1 mL, 2 times), fixed with 4% PFA (0.5 mL) for 20 min at RT, and washed with PBS (1 mL, 2 times). The cells were permeabilized in PBS supplemented with 0.1% Triton X-100 (0.5 mL) for 5 min, successively stained with Alexa Fluor 488-phalloidin and DAPI (1 mL) according to the manufacturer’s protocols and imaged by confocal microscopy using a Leica TCS SP5 X WLL instrument. Tile images were recreated from five individual O_2_-cryogel samples to visualize all the dimensions of the gel, and cell viability was calculated by Fiji software(*88*) as the ratio between the dead cell number and the total cell number, which correspond to surface areas positive for far-red and blue or green fluorescence, respectively. For each condition, one representative high-resolution stacked image was collected with a 2-μm separation between slices (z-stacks).

### Minimally invasive delivery and biocompatibility assessment

Seven-week-old female C57BL/6J mice (n = 4; The Jackson Laboratory, Bar Harbor, Maine, USA) were anesthetized with isoflurane (4–5% for induction and 1–3% for maintenance) in oxygen using an inhalation anesthesia system (300 SERIES vaporizer, VSS, Rockmart, GA, USA). Cryogels (i.e., CaO_2_-free cryogels) and O_2_-cryogels (i.e., CaO_2_-containing cryogels) were suspended in PBS (0.2 mL) and subcutaneously injected into both dorsal flanks of each mouse using a 16-gauge hypodermic needle. After 7 days, mice were euthanized, and cryogels and O_2_-cryogels were explanted from the surrounding tissues. Next, each sample was fixed for 48 h in 4% PFA (2 mL), embedded in paraffin, cryosectioned into 5-μm-thick slices, and stained with hematoxylin and eosin (H&E) as well as Masson’s trichrome (MT) for histological analysis.

### Hypoxia detection assays

Cellular hypoxia within O_2_-cryogels was assessed using a Hypoxyprobe™-1 kit according to the manufacturer’s recommendations. Briefly, after 24 h of incubation in normoxia or hypoxia, cell-laden O_2_-cryogels were incubated for 2 h in 200 µM Hypoxyprobe™-1 in complete DMEM (2 mL), fixed, permeabilized, and immunostained with mouse MAb1 antibody and rabbit anti-mouse IgG (H+L) Alexa Fluor 488 secondary antibody, as well as DAPI. Cryogel samples (cryogels and O_2_-cryogels) were then imaged by confocal microscopy using the Leica TCS SP5 X WLL instrument. Cellular hypoxia fractions were calculated using Fiji software(*88*) based on the ratio between the number of hypoxic cells (green fluorescence) and the total number of cells (blue fluorescence).

Adenosine concentration was quantified using a fluorometric adenosine assay kit following the manufacturer’s protocol. After 24 h of incubation under either normoxic or hypoxic conditions, supernatants from each cell-laden cryogel samples (cryogels and O_2_-cryogels) were collected and mixed with the assay kit. The adenosine concentration was determined based on the fluorescence intensity (Ex/Em = 535 nm/587 nm), which was measured using a Synergy HT plate reader (BioTek) and normalized using an adenosine standard curve.

The gene expression of HIF1α, wingless-type MMTV integration site family member 11 (Wnt11), vascular endothelial growth factor α (VEGFα), cluster of differentiation 44 (CD44), CD73, CD133, and hypoxanthine guanine phosphoribosyl transferase (HPRT) were assessed by RT-qPCR. After 24 h of incubation in either normoxia or hypoxia, total RNA was extracted from cell-laden cryogel samples using a PureLink™ RNA Mini Kit according to the manufacturer’s protocol. For RT-qPCR, we first used a High Capacity cDNA Reverse Transcription Kit on a MyCycler (Bio-Rad, Hampton, NH, USA). Then, gene expression was measured using the following TaqMan® Gene Expression Assays on an Mx3005P QPCR System (Agilent, Santa Clara, CA, USA): *HIF1α*: Mm00468869_m1, *Wnt11*: Mm00437327_g1, *VEGFα*: Mm00437306_m1, *CD44*: Mm01277163_m1, *CD73*: Mm00501915_m1, *CD133*: Mm00477115_m1, and *HPRT* (housekeeping gene): Mm03024075_m1.

### Cytotoxicity assay in a B16-OVA melanoma model

B16-OVA cells were stained with 10 μM CFSE for 15 min at 37 °C according to the manufacturer’s protocol. Next, 1×10^5^ cells in complete Roswell Park Memorial Institute (RPMI) medium (RPMI medium supplemented with 10% FBS, 100 µg/mL penicillin, and 100 µg/mL streptomycin) were seeded onto square-shaped cryogels and incubated in normoxia. After 3 h of incubation, splenocytes isolated from OT-1 RAG mice (The Jackson Laboratory, Bar Harbor, ME, USA) in complete RPMI (2×10^5^ cells, 1:2 target/effector ratio) were added to each well. Splenocytes were prestimulated with 2μg/mL SIINFEKL peptides for 24 h in complete RPMI supplemented with 50 µM 2-mercaptoethanol. For experiments requiring a hypoxic environment, complete RPMI, splenocyte suspensions, and B16-OVA-laden O_2_-cryogels were preincubated (conditioned) for 1 h in hypoxia prior to coculture. After 24 h of B16-OVA/splenocyte coculture, (*i*) supernatants were collected to quantify IFNγ, TNFα, granzyme B, and perforin secretion by ELISA; and (*ii*) the cytotoxicity of OT-1 cells was assessed by the cell viability assay described above.

The cytolytic activity of OT-1 T cells towards B16-OVA cells was monitored by time-lapse microscopy. First, under normoxic conditions, 1×10^5^ B16-OVA cells in complete RPMI were seeded in a 24-well plate for 3 h to allow adhesion. Next, 2×10^5^ activated splenocytes in complete RPMI (1:2 target/effector ratio) with either square-shaped cryogels or O_2_-cryogels containing 0.5% CaO_2_ were added to each well (total volume = 500 µL). Complete RPMI, splenocyte suspensions, and the 24-well plate seeded with B16-OVA cells were preincubated (conditioned) for 1 h in hypoxia prior to coculture. Next, time-lapse experiments were performed in a closed chamber using a bright field inverted microscope (Zeiss Axio Observer Z1, Zeiss, Oberkochen, Germany) at 37 °C under hypoxic conditions (5% CO_2_ + 1% O_2_ + 94% N_2_). One preselected region of each well (350 × 350 µm) was imaged every 10 min for 4 h.

### Intratumoral oxygen delivery as a therapeutic co-adjuvant in advanced-stage tumor-bearing mice

Murine melanoma tumors (two tumors/mouse) were established in C57BL/6 mice by subcutaneous injection of B16-F10 cells (2×10^5^ cells in 100 µL PBS) in both flanks. Seventeen days following tumor inoculation (tumor volume > 150 mm^3^), mice were treated twice a week with either intratumoral injection of PBS (100 µL, sham), cylindrical-shaped 1% O_2_-cryogel (height: 35 mm, diameter: 1.2 mm; in 100 µL PBS) or placed in chambers with controlled oxygen content (60%) to mimic supplemental oxygen therapy to humans.(*89*) Tumor volume was measured twice a week using a caliper. Once tumors exceeded 1 cm^3^, or severe ulceration, bleeding, or weight variation were observed, mice were euthanized. To assess the antitumor immune response, cryogels, spleens, and lymph nodes (LNs) were explanted and dissociated at day 24, and the immune cell populations were analyzed by flow cytometry (AttuneNxT, Thermo Fisher Scientific). The following markers were used to determine the T cell, NK cell, and hypoxic cell subpopulations: CD45 (immune cells), Hypoxyprobe™-1 (hypoxic cells), NK1.1 (natural killer cells), PD-1 (checkpoint inhibitor), CD3 (T cells), CD4 (T helper cells), CD8 (cytotoxic T cells), CD25 and FoxP3 (regulatory T cells). To evaluate the systemic inflammatory response, mouse blood was collected at day 24 (50 µL), and the level of IL-2, IL-4, IL-5, IL-6, IL-9, IL-13, IL-17A, IL-17F, IL-22, IFN-γ, and TNF-α in serum was analyzed using LEGENDplex Mouse Th Cytokine Panel.

### Animal studies

All experiments requiring mice were conducted in compliance with the National Institutes of Health (NIH) guidelines and approved by the Division of Laboratory Animal Medicine and Northeastern University Institutional Animal Care and Use Committee (protocol number 17-0828R).

### Statistical analysis

All data are presented as the mean ± standard error of the mean (SEM). Statistical analyses were performed using GraphPad Prism software (La Jolla, CA, USA). Significant differences between groups were analyzed by one-way analysis of variance (ANOVA) and Dunnett’s or Bonferroni’s post-hoc test, two-tailed Pearson test, or a two-tailed Mantel-Cox text. Survival analysis was performed with the Kaplan-Meier test. As a proxy for life expectancy, the median survival time was determined as the time when a survival curve reaches 50% on the y-axis. Differences were considered statistically significant at P < 0.05.

## Acknowledgements

S.A.B. acknowledges the support from Northeastern University Seed Grant/Proof of Concept Tier 1 Research Grant, Burroughs Wellcome Fund award (1018798), Thomas Jefferson/Face foundation award, DFCI/NU Joint Program Grant, and NSF CAREER award (DMR 1847843). S.A.B. and A.M. acknowledge the support from the National Plan for Science, Technology and Innovation (MAARIFAH), King Abdulaziz City for Science and Technology, the Kingdom of Saudi Arabia, award (15-MED5025-03). S.A.B. and A.M. also acknowledge with thanks the Science and Technology Unit, King Abdulaziz University for technical support. R. Koppes, A. Koppes, and J. Soucy are acknowledged for their technical support with time-lapse microscopy. Thanks to Edward Emmott, Gray Huffman, and Zachary Rogers for proofreading the manuscript.

## Author contributions

T.C. and S.A.B. conceived and designed the experiments. T.C., L.J.E, and M.R. performed the experiments. T.C., S.H., L.J.E., A.M., M.V.S and S.A.B. analyzed the data and wrote the manuscript. T.C., L.J.E., A.M. and S.A.B. conceived the figures. All authors discussed the results and commented the manuscript. The principal investigator is S.A.B.

## References

1. K. Esfahani et al., A review of cancer immunotherapy: from the past, to the present, to the future. Curr Oncol 27, S87–S97 (2020).

2. M. Sitkovsky, D. Lukashev, Regulation of immune cells by local-tissue oxygen tension: HIF1 alpha and adenosine receptors. Nature reviews. Immunology 5, 712–721 (2005).

3. M. V. Sitkovsky et al., Hostile, hypoxia-A2-adenosinergic tumor biology as the next barrier to overcome for tumor immunologists. Cancer Immunol Res 2, 598–605 (2014).

4. D. N. Khalil, E. L. Smith, R. J. Brentjens, J. D. Wolchok, The future of cancer treatment: immunomodulation, CARs and combination immunotherapy. Nature Reviews Clinical Oncology 13, 273–290 (2016).

5. A. Arina, L. Corrales, V. Bronte, Enhancing T cell therapy by overcoming the immunosuppressive tumor microenvironment. Seminars in Immunology 28, 54–63 (2016).

6. W. Al Tameemi, T. P. Dale, R. M. K. Al-Jumaily, N. R. Forsyth, Hypoxia-Modified Cancer Cell Metabolism. Frontiers in Cell and Developmental Biology 7, (2019).

7. P. Vaupel, A. Mayer, Hypoxia in cancer: significance and impact on clinical outcome. Cancer metastasis reviews 26, 225–239 (2007).

8. E. Paolicchi et al., Targeting hypoxic response for cancer therapy. Oncotarget 7, 13464–13478 (2016).

9. M. R. Horsman, J. Overgaard, The impact of hypoxia and its modification of the outcome of radiotherapy. Journal of Radiation Research 57, i90–i98 (2016).

10. S. Rey, L. Schito, M. Koritzinsky, B. G. Wouters, Molecular targeting of hypoxia in radiotherapy. Advanced Drug Delivery Reviews 109, 45–62 (2017).

11. A. Sermeus et al., Hypoxia-Induced Modulation of Apoptosis and BCL-2 Family Proteins in Different Cancer Cell Types. PLOS ONE 7, e47519 (2012).

12. B. Muz, P. de la Puente, F. Azab, A. K. Azab, The role of hypoxia in cancer progression, angiogenesis, metastasis, and resistance to therapy. Hypoxia 3, 83–92 (2015).

13. E. B. Rankin, A. J. Giaccia, Hypoxic control of metastasis. Science (New York, N.Y.) 352, 175–180 (2016).

14. A. Ohta et al., A2A adenosine receptor protects tumors from antitumor T cells. Proceedings of the National Academy of Sciences 103, 13132–13137 (2006).

15. B. Deng et al., Intratumor Hypoxia Promotes Immune Tolerance by Inducing Regulatory T Cells via TGF-β1 in Gastric Cancer. PLOS ONE 8, e63777 (2013).

16. S. M. Hatfield et al., Systemic oxygenation weakens the hypoxia and Hypoxia Inducible Factor 1α-dependent and extracellular adenosine-mediated tumor protection. Journal of molecular medicine (Berlin, Germany) 92, 1283–1292 (2014).

17. S. M. Hatfield et al., Immunological mechanisms of the antitumor effects of supplemental oxygenation. Science translational medicine 7, 277ra230 (2015).

18. G. L. Semenza, G. L. Wang, A nuclear factor induced by hypoxia via de novo protein synthesis binds to the human erythropoietin gene enhancer at a site required for transcriptional activation. Molecular and Cellular Biology 12, 5447–5454 (1992).

19. H. Kojima et al., Abnormal B lymphocyte development and autoimmunity in hypoxia-inducible factor 1alpha-deficient chimeric mice. Proceedings of the National Academy of Sciences of the United States of America 99, 2170–2174 (2002).

20. M. Thiel et al., Targeted deletion of HIF-1alpha gene in T cells prevents their inhibition in hypoxic inflamed tissues and improves septic mice survival. PLoS One 2, e853 (2007).

21. M. V. Sitkovsky et al., Hostile, Hypoxia-A2-Adenosinergic Tumor Biology as the Next Barrier to the Tumor Immunologists. Cancer immunology research 2, 598–605 (2014).

22. S. M. Hatfield, M. Sitkovsky, Oxygenation to improve cancer vaccines, adoptive cell transfer and blockade of immunological negative regulators. Oncoimmunology 4, e1052934 (2015).

23. A. M. Graham, K. G. McCracken, Convergent evolution on the hypoxia-inducible factor (HIF) pathway genes EGLN1 and EPAS1 in high-altitude ducks. Heredity 122, 819–832 (2019).

24. S. M. Hatfield et al., Systemic oxygenation weakens the hypoxia and hypoxia inducible factor 1alpha-dependent and extracellular adenosine-mediated tumor protection. Journal of molecular medicine (Berlin, Germany) 92, 1283–1292 (2014).

25. V. Simko et al., Hypoxia induces cancer-associated cAMP/PKA signalling through HIF-mediated transcriptional control of adenylyl cyclases VI and VII. Scientific Reports 7, 10121 (2017).

26. M. Sitkovsky, D. Lukashev, Regulation of immune cells by local-tissue oxygen tension: HIF1α and adenosine receptors. Nature Reviews Immunology 5, 712 (2005).

27. M. V. Sitkovsky, J. Kjaergaard, D. Lukashev, A. Ohta, Hypoxia-Adenosinergic Immunosuppression: Tumor Protection by T Regulatory Cells and Cancerous Tissue Hypoxia. Clinical Cancer Research 14, 5947–5952 (2008).

28. E. N. McNamee, D. Korns Johnson, D. Homann, E. T. Clambey, Hypoxia and hypoxia-inducible factors as regulators of T cell development, differentiation, and function. Immunologic research 55, 58–70 (2013).

29. S. Sarkar et al., Hypoxia Induced Impairment of NK Cell Cytotoxicity against Multiple Myeloma Can Be Overcome by IL-2 Activation of the NK Cells. PLOS ONE 8, e64835 (2013).

30. C. Wigerup, S. Pahlman, D. Bexell, Therapeutic targeting of hypoxia and hypoxia-inducible factors in cancer. Pharmacology & therapeutics 164, 152–169 (2016).

31. S. M. Hatfield, M. Sitkovsky, A2A Adenosine Receptor antagonists to weaken the hypoxia-HIF-1α driven immunosuppression and improve immunotherapies of cancer. Current opinion in pharmacology 29, 90–96 (2016).

32. J. Kjaergaard, S. Hatfield, G. Jones, A. Ohta, M. Sitkovsky, A2A Adenosine Receptor Gene Deletion or Synthetic A2A Antagonist Liberate Tumor-Reactive CD8(+) T Cells from Tumor-Induced Immunosuppression. Journal of immunology (Baltimore, Md. : 1950) 201, 782–791 (2018).

33. S. K. Burroughs et al., Hypoxia inducible factor pathway inhibitors as anticancer therapeutics. Future medicinal chemistry 5, 10.4155/fmc.4113.4117 (2013).

34. I. Moen, L. E. B. Stuhr, Hyperbaric oxygen therapy and cancer—a review. Targeted Oncology 7, 233–242 (2012).

35. R. D. Leone, L. A. Emens, Targeting adenosine for cancer immunotherapy. Journal for ImmunoTherapy of Cancer 6, 57 (2018).

36. B. R. O’Driscoll, L. S. Howard, A. G. Davison, BTS guideline for emergency oxygen use in adult patients. Thorax 63, vi1–vi68 (2008).

37. L. Gu, D. J. Mooney, Biomaterials and emerging anticancer therapeutics: engineering the microenvironment. Nature Reviews Cancer 16, 56 (2015).

38. J. S. Weber, J. J. Mulé, Cancer immunotherapy meets biomaterials. Nature Biotechnology 33, 44 (2015).

39. G. Camci-Unal, N. Alemdar, N. Annabi, A. Khademhosseini, Oxygen Releasing Biomaterials for Tissue Engineering. Polymer international 62, 843–848 (2013).

40. A. L. Farris, A. N. Rindone, W. L. Grayson, Oxygen Delivering Biomaterials for Tissue Engineering. Journal of materials chemistry. B, Materials for biology and medicine 4, 3422–3432 (2016).

41. N. Alemdar et al., Oxygen-Generating Photo-Cross-Linkable Hydrogels Support Cardiac Progenitor Cell Survival by Reducing Hypoxia-Induced Necrosis. ACS Biomaterials Science & Engineering 3, 1964–1971 (2017).

42. B. Newland, M. Baeger, D. Eigel, H. Newland, C. Werner, Oxygen-Producing Gellan Gum Hydrogels for Dual Delivery of Either Oxygen or Peroxide with Doxorubicin. ACS Biomaterials Science & Engineering 3, 787–792 (2017).

43. M. Gholipourmalekabadi, S. Zhao, B. S. Harrison, M. Mozafari, A. M. Seifalian, Oxygen-Generating Biomaterials: A New, Viable Paradigm for Tissue Engineering? Trends in biotechnology 34, 1010–1021 (2016).

44. W. A. Li, D. J. Mooney, Materials based tumor immunotherapy vaccines. Current Opinion in Immunology 25, 238–245 (2013).

45. S. Park, K. M. Park, Hyperbaric oxygen-generating hydrogels. Biomaterials 182, 234–244 (2018).

46. P. A. Shiekh, A. Singh, A. Kumar, Oxygen-Releasing Antioxidant Cryogel Scaffolds with Sustained Oxygen Delivery for Tissue Engineering Applications. ACS applied materials & interfaces 10, 18458–18469 (2018).

47. F. Dehghani, N. Annabi, Engineering porous scaffolds using gas-based techniques. Current Opinion in Biotechnology 22, 661–666 (2011).

48. K. J. De France, F. Xu, T. Hoare, Structured Macroporous Hydrogels: Progress, Challenges, and Opportunities. Advanced healthcare materials 7, (2018).

49. S. A. Bencherif et al., Injectable preformed scaffolds with shape-memory properties. Proceedings of the National Academy of Sciences 109, 19590–19595 (2012).

50. A. Memic et al., Latest Advances in Cryogel Technology for Biomedical Applications. Advanced Therapeutics 2, 1800114 (2019).

51. F. A. Venning, L. Wullkopf, J. T. Erler, Targeting ECM Disrupts Cancer Progression. Frontiers in Oncology 5, 224 (2015).

52. M. J. Cooke, K. Vulic, M. S. Shoichet, Design of biomaterials to enhance stem cell survival when transplanted into the damaged central nervous system. Soft Matter 6, 4988–4998 (2010).

53. M. Gülden, A. Jess, J. Kammann, E. Maser, H. Seibert, Cytotoxic potency of H2O2 in cell cultures: Impact of cell concentration and exposure time. Free Radical Biology and Medicine 49, 1298–1305 (2010).

54. Y. Li, S. P. Patel, J. Roszik, Y. Qin, Hypoxia-Driven Immunosuppressive Metabolites in the Tumor Microenvironment: New Approaches for Combinational Immunotherapy. Front Immunol 9, 1591–1591 (2018).

55. P. Bhat, G. Leggatt, N. Waterhouse, I. H. Frazer, Interferon-γ derived from cytotoxic lymphocytes directly enhances their motility and cytotoxicity. Cell Death & Disease 8, e2836–e2836 (2017).

56. B. Lowin, M. Hahne, C. Mattmann, J. Tschopp, Cytolytic T-cell cytotoxicity is mediated through perforin and Fas lytic pathways. Nature 370, 650–652 (1994).

57. C. M. L. West et al., Targeting Hypoxia to Improve Non–Small Cell Lung Cancer Outcome.JNCI: Journal of the National Cancer Institute 110, 14–30 (2017).

58. L. Figueiredo et al., Assessing glucose and oxygen diffusion in hydrogels for the rational design of 3D stem cell scaffolds in regenerative medicine. Journal of Tissue Engineering and Regenerative Medicine 12, 1238–1246 (2018).

59. S. H. Oh, C. L. Ward, A. Atala, J. J. Yoo, B. S. Harrison, Oxygen generating scaffolds for enhancing engineered tissue survival. Biomaterials 30, 757–762 (2009).

60. C.-S. Yeh, R. Wang, W.-C. Chang, Y.-h. Shih, Synthesis and characterization of stabilized oxygen-releasing CaO2 nanoparticles for bioremediation. Journal of Environmental Management 212, 17–22 (2018).

61. E. Pedraza, M. M. Coronel, C. A. Fraker, C. Ricordi, C. L. Stabler, Preventing hypoxia-induced cell death in beta cells and islets via hydrolytically activated, oxygen-generating biomaterials. Proceedings of the National Academy of Sciences of the United States of America 109, 4245–4250 (2012).

62. S. I. H. Abdi, S. M. Ng, J. O. Lim, An enzyme-modulated oxygen-producing micro-system for regenerative therapeutics. International journal of pharmaceutics 409, 203–205 (2011).

63. S. F. Khattak, K. S. Chin, S. R. Bhatia, S. C. Roberts, Enhancing oxygen tension and cellular function in alginate cell encapsulation devices through the use of perfluorocarbons. Biotechnology and bioengineering 96, 156–166 (2007).

64. M. A. Mensink, H. W. Frijlink, K. van der Voort Maarschalk, W. L. J. Hinrichs, How sugars protect proteins in the solid state and during drying (review): Mechanisms of stabilization in relation to stress conditions. European Journal of Pharmaceutics and Biopharmaceutics 114, 288–295 (2017).

65. C. W. Pugh, P. J. Ratcliffe, Regulation of angiogenesis by hypoxia: role of the HIF system. Nature Medicine 9, 677–684 (2003).

66. S. Chouaib et al., Hypoxia Promotes Tumor Growth in Linking Angiogenesis to Immune Escape. Frontiers in Immunology 3, (2012).

67. H. Mori et al., Induction of WNT11 by hypoxia and hypoxia-inducible factor-1α regulates cell proliferation, migration and invasion. Scientific Reports 6, 21520 (2016).

68. E. Monzani et al., Melanoma contains CD133 and ABCG2 positive cells with enhanced tumourigenic potential. European Journal of Cancer 43, 935–946 (2007).

69. A. Dietrich, E. Tanczos, W. Vanscheidt, E. Schöpf, J. C. Simon, High CD44 surface expression on primary tumours of malignant melanoma correlates with increased metastatic risk and reduced survival. European Journal of Cancer 33, 926–930 (1997).

70. L. L. van der Woude, M. A. J. Gorris, A. Halilovic, C. G. Figdor, I. J. M. de Vries, Migrating into the Tumor: a Roadmap for T Cells. Trends in Cancer 3, 797–808 (2017).

71. S. Sethumadhavan et al., Hypoxia and hypoxia-inducible factor (HIF) downregulate antigen-presenting MHC class I molecules limiting tumor cell recognition by T cells. PLoS One 12, e0187314 (2017).

72. K. Stępień, R. P. Ostrowski, E. Matyja, Hyperbaric oxygen as an adjunctive therapy in treatment of malignancies, including brain tumours. Med Oncol 33, 101–101 (2016).

73. J. F. Turrens, Mitochondrial formation of reactive oxygen species. The Journal of physiology 552, 335–344 (2003).

74. M. Z. Noman et al., Hypoxia: a key player in antitumor immune response. A Review in the Theme: Cellular Responses to Hypoxia. Am J Physiol Cell Physiol 309, C569–C579 (2015).

75. E. Maniati, R. Soper, T. Hagemann, Up for Mischief? IL-17/Th17 in the tumour microenvironment. Oncogene 29, 5653–5662 (2010).

76. H. Okamoto et al., Interleukin-13 receptor α2 is a novel marker and potential therapeutic target for human melanoma. Scientific Reports 9, 1281 (2019).

77. N. Shimasaki, A. Jain, D. Campana, NK cells for cancer immunotherapy. Nature Reviews Drug Discovery 19, 200–218 (2020).

78. T. Y. Shih et al., Injectable, Tough Alginate Cryogels as Cancer Vaccines. Advanced healthcare materials 7, e1701469 (2018).

79. S. A. Bencherif et al., Injectable cryogel-based whole-cell cancer vaccines. Nature Communications 6, 7556 (2015).

80. T. Colombani et al., Lipidic Aminoglycoside Derivatives: A New Class of Immunomodulators Inducing a Potent Innate Immune Stimulation. Adv Sci (Weinh) 6, 1900288–1900288 (2019).

81. J. Kim et al., Injectable, spontaneously assembling, inorganic scaffolds modulate immune cells in vivo and increase vaccine efficacy. Nature biotechnology 33, 64–72 (2015).

82. J. W. Hickey et al., Immunotherapy: Engineering an Artificial T-Cell Stimulating Matrix for Immunotherapy (Adv. Mater. 23/2019). Advanced Materials 31, 1970162 (2019).

83. M. Mahmoudi, O. C. Farokhzad, Cancer immunotherapy: Wound-bound checkpoint blockade. Nature Biomedical Engineering 1, 0031 (2017).

84. V. Petrova, M. Annicchiarico-Petruzzelli, G. Melino, I. Amelio, The hypoxic tumour microenvironment. Oncogenesis 7, 10 (2018).

85. P. Jayaprakash et al., Targeted hypoxia reduction restores T cell infiltration and sensitizes prostate cancer to immunotherapy. J Clin Invest 128, 5137–5149 (2018).

86. R. D. Leone, Y.-C. Lo, J. D. Powell, A2aR antagonists: Next generation checkpoint blockade for cancer immunotherapy. Computational and Structural Biotechnology Journal 13, 265–272 (2015).

87. M. Rezaeeyazdi, T. Colombani, A. Memic, S. A. Bencherif, Injectable Hyaluronic Acid-co-Gelatin Cryogels for Tissue-Engineering Applications. Materials (Basel, Switzerland) 11, (2018).

88. J. Schindelin et al., Fiji: an open-source platform for biological-image analysis. Nature Methods 9, 676 (2012).

89. M. Thiel et al., Oxygenation inhibits the physiological tissue-protecting mechanism and thereby exacerbates acute inflammatory lung injury. PLoS Biol 3, e174–e174 (2005).

