## Supplementary Figure 1 for "Oxygen-generating cryogels restore T cell-mediated cytotoxicity in hypoxic tumors"

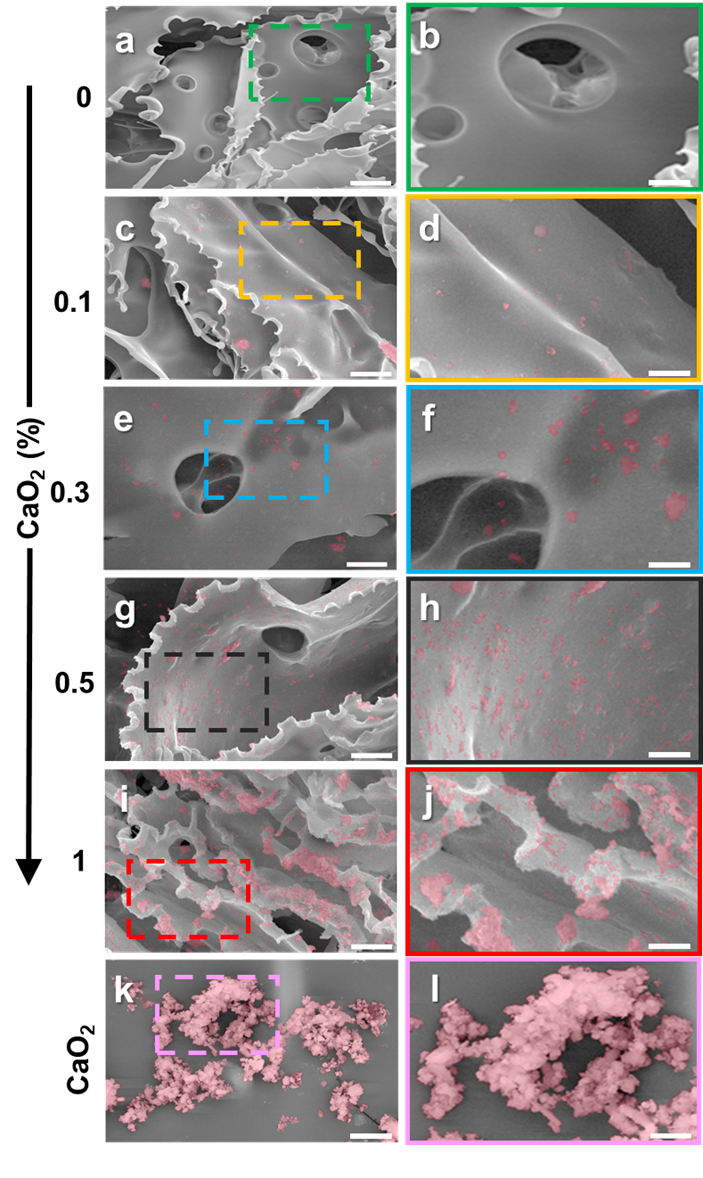

**Supplementary Figure 1. Scanning electron microscopy images of entrapped CaO_2_ particles within the polymer walls of O_2_-cryogels.** Representative images (n = 5) showing various concentrations of CaO_2_ particles within the polymer walls. (a,b) cryogels (0% CaO_2_, control), O_2_-cryogels containing (c, d) 0.1% CaO_2_,(e, f) 0.3% CaO_2_, (g, h) 0.5% CaO_2_, and (i, j) 1% CaO_2_, and free CaO_2_ particles (k, l; control). Pseudocoloring in pink is used to highlight CaO_2_ particles. Scale bars = 50 µm (a, c, e, g, i), 25 µm (b, d, f, h, j), 10 µm (k), and 5 µm (l).

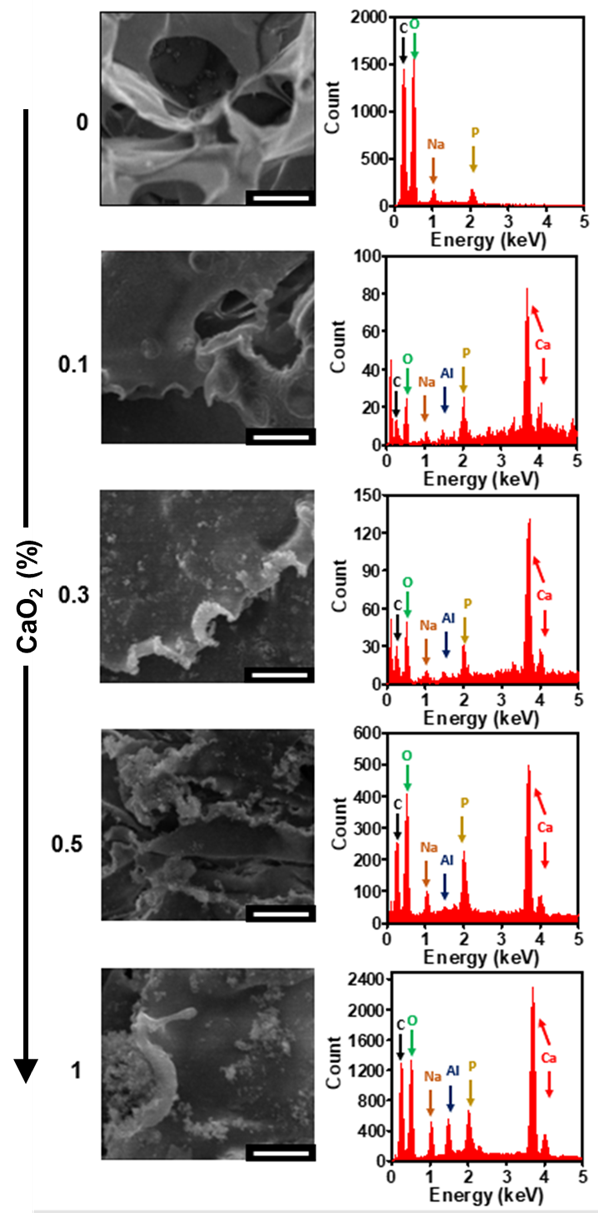

**Supplementary Figure 2. Energy-dispersive X-ray spectroscopy (EDX) microanalysis of CaO_2_-containing cryogels.** Elemental analysis of cryogels and O_2_-cryogels prepared with 0.1%, 0.3%, 0.5%, and 1% (wt/v) CaO_2_ particles. Scanning electron microscopy images (left) of the area analyzed by EDX (right). The data are representative of n = 5 cryogels per condition. Scale bars = 25 µm.

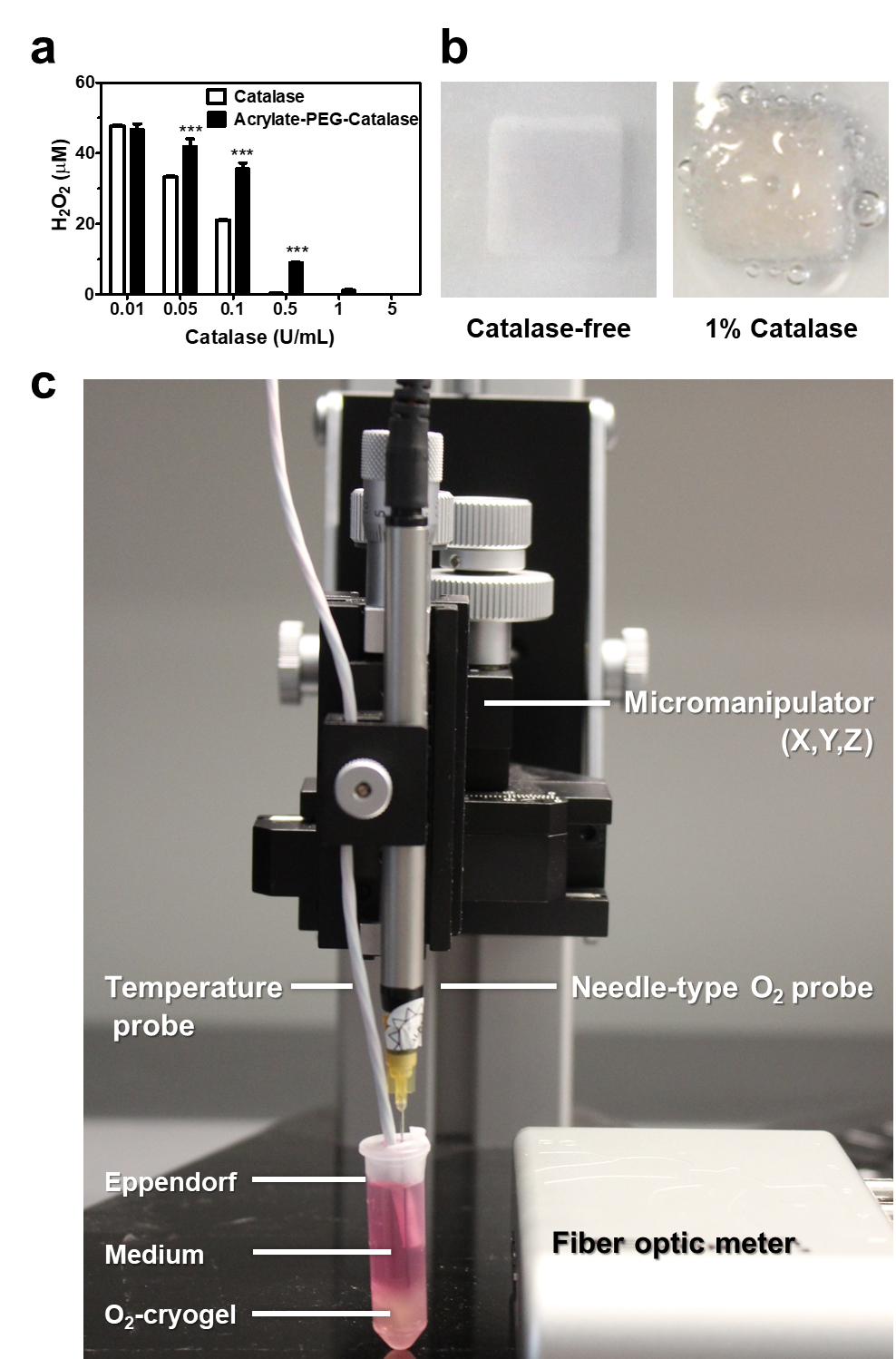

**Supplementary Figure 3. Catalase-mediated H_2_O_2_ depletion and cryogel-mediated oxygen release.** a) Enzymatic activity of catalase (nonconjugated) and APC in 50 μM H_2_O_2_. b) Photographs of catalase-free (left) and 1% wt/v catalase-containing cryogels (right, active oxygen release produces bubbles) immersed in a 50 mM H_2_O_2_ solution for 5 s. c) Experimental setup. The values represent the mean ± SEM (n = 6). Data were analyzed using ANOVA and Dunnett’s post hoc test (compared to nonconjugated catalase), ^***^P < 0.001.

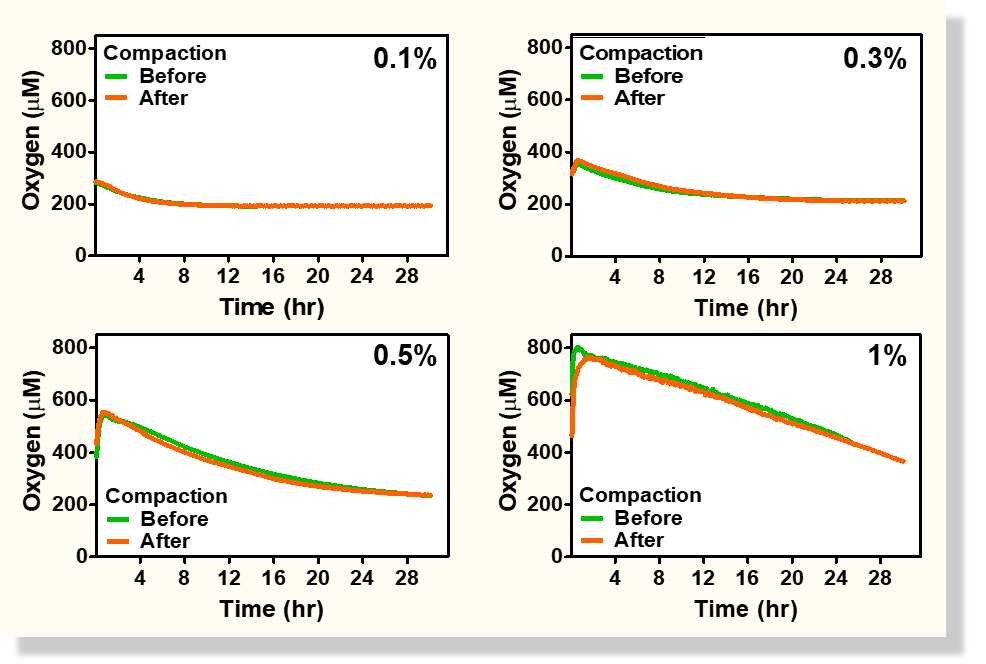

**Supplementary Figure 4. Controlled oxygen release from O_2_-cryogels upon injection-like conditions.** Oxygen release from O_2_-cryogels containing various concentration of CaO_2_ (0.1-1%) before (green) and after mechanical compaction (orange) under normoxic conditions. Values represent the mean ± SEM (n = 4).

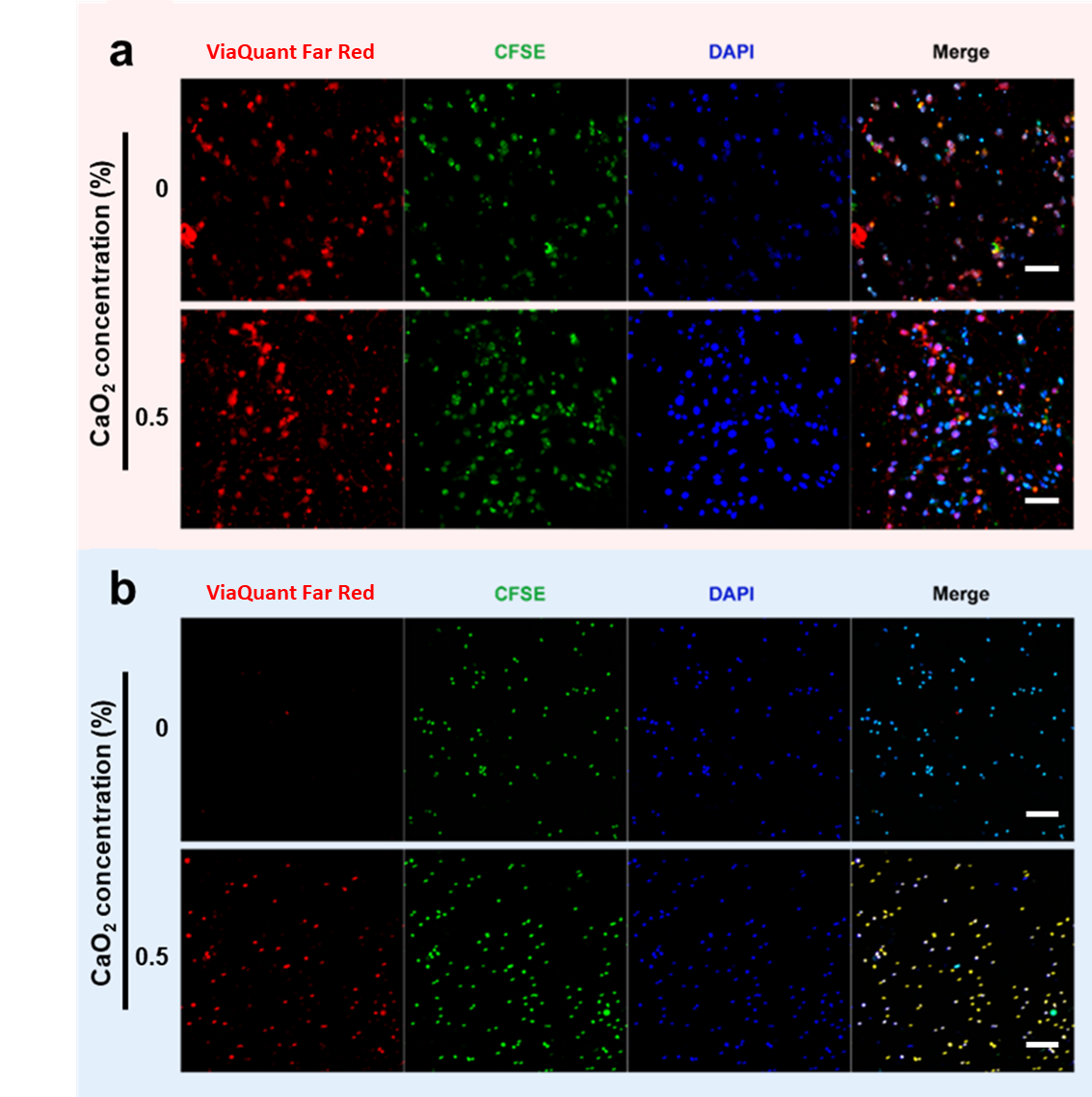

**Supplementary Figure 5.** **Cytotoxic activity of OT-1 T cells cultured in B16-OVA cell-laden O_2_-cryogels.** a-b) Confocal images depicting B16-OVA cell viability within O_2_-cryogels after 24 h of incubation with OT-1 T cells under normoxic (20% O_2_) (a) or hypoxic conditions (1% O_2_) (b). Red = dead cells stained with ViaQuant Far Red, blue = nuclei stained with DAPI, green = B16-OVA cells stained with CFSE. The data are representative of n = 5 samples per condition. Scale bars = 50 µm (a) and 100 µm (b).

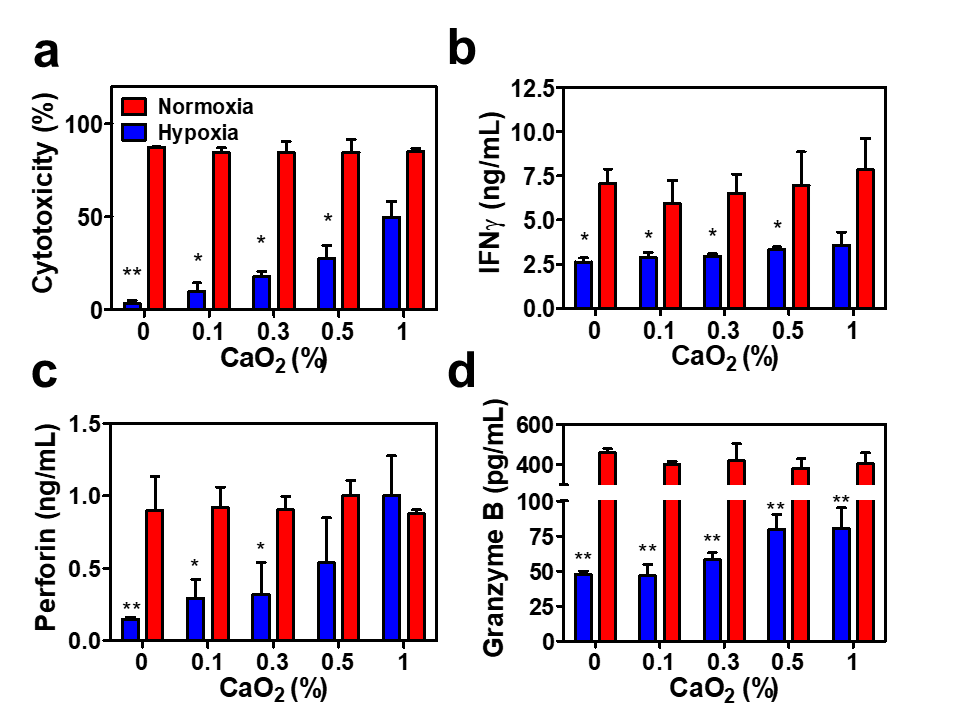

**Supplementary Figure 6. Rescuing the cytotoxic activity of hypoxic T cells with O_2_-cryogels.** a-d) B16-OVA cells were cocultured with OT-1 T cells for 4 h under normoxic (20% O_2_, red) or hypoxic conditions (1% O_2_, blue). a) Quantification of T cell-mediated cytotoxicity against B16-OVA cells. b-d) Secretion of IFNγ, (b) perforin (c), and granzyme B (d) from OT-1 T cells. Values represent the mean ± SEM (n = 4). Data were analyzed using ANOVA and Dunnett’s post hoc test (compared to cryogels), ^*^P < 0.5, ^**^P < 0.01.

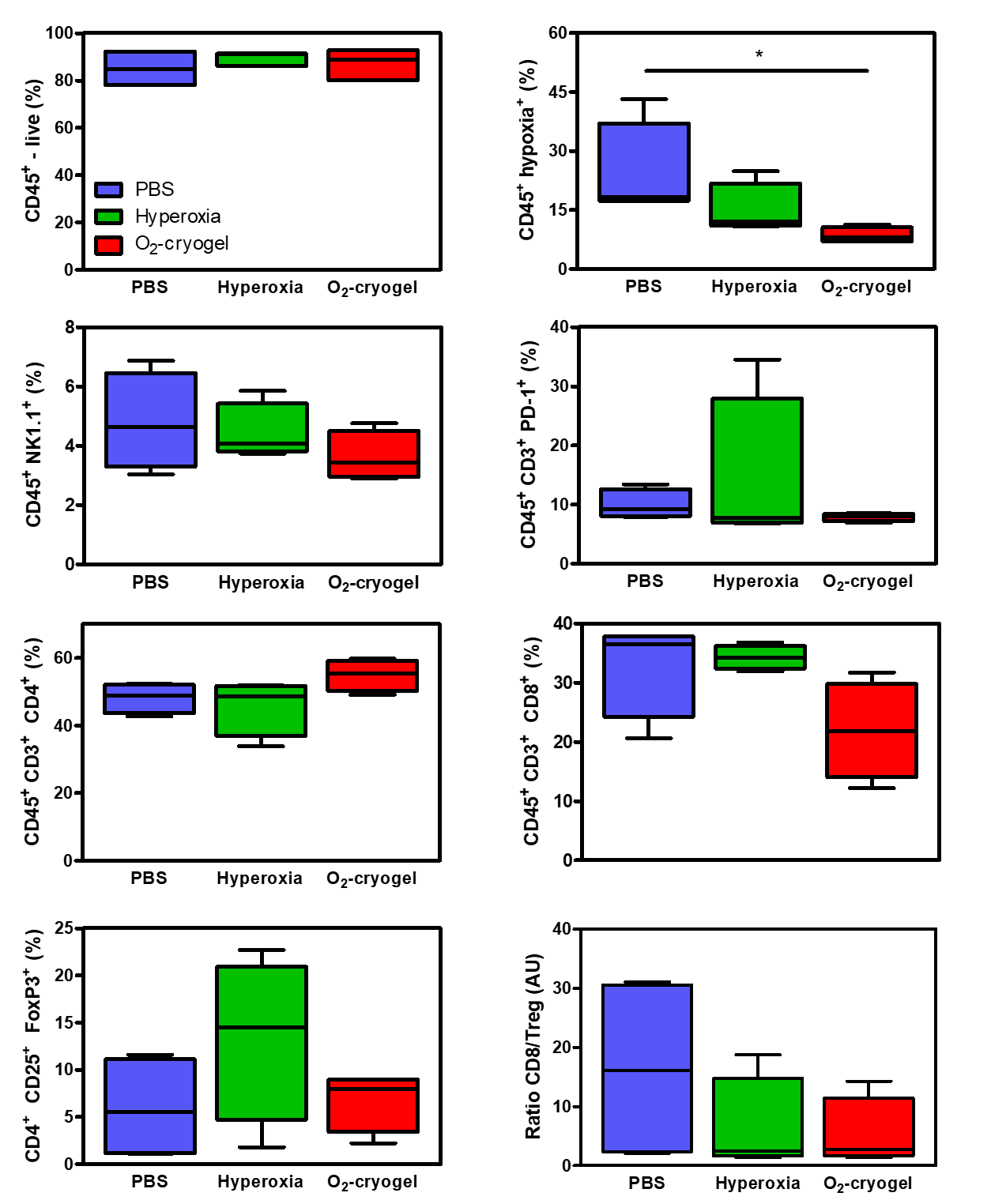

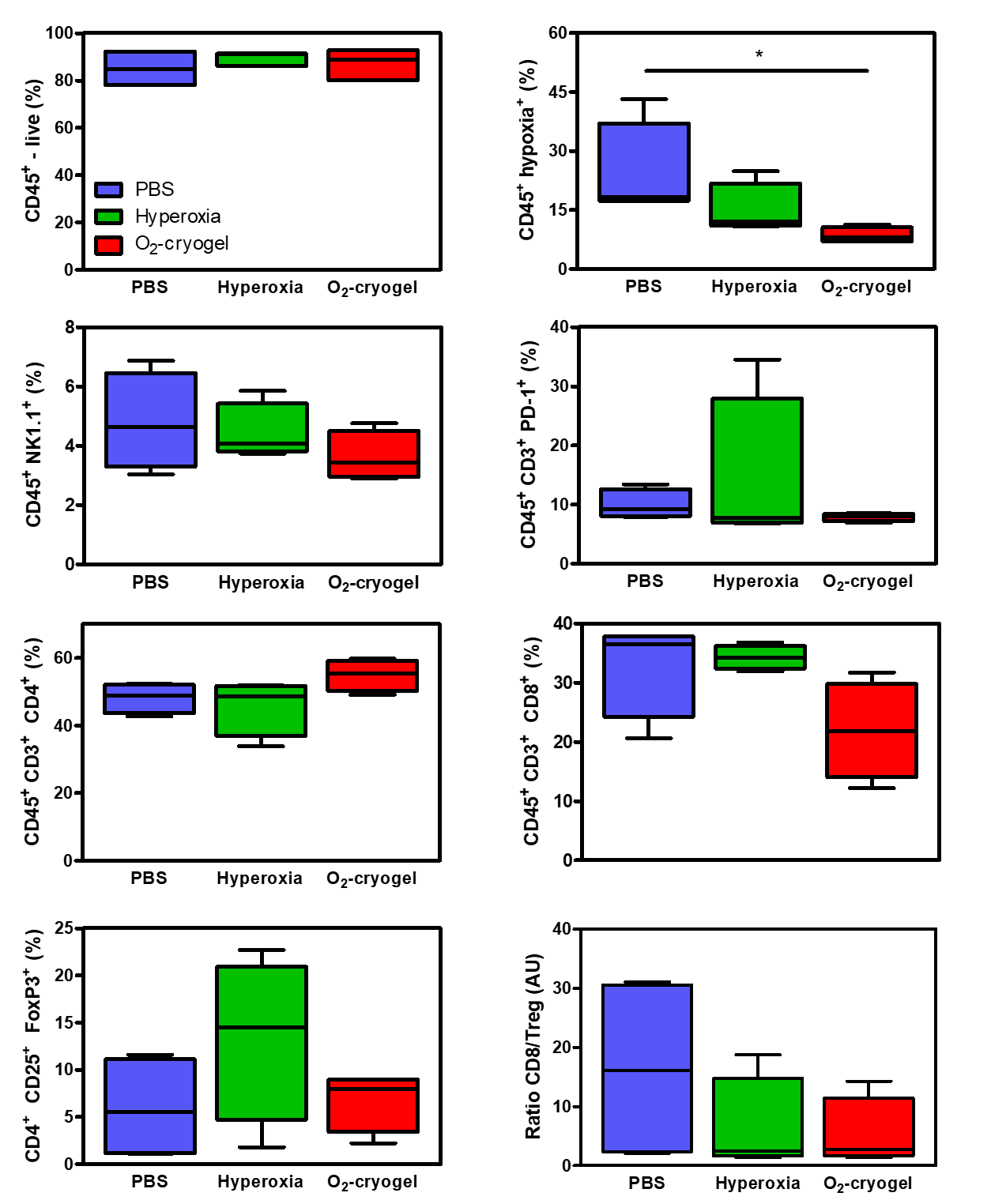

**Supplementary Figure 7. Changes in the frequency of immune cell population following O_2_-cryogel co-adjuvant therapy.** Fractions of immune cells (CD45^+^), hypoxic immune cells (CD45^+^ Hypoxia^+^), natural killer cells (CD45^+^ NK1.1^+^), PD-1 positive T cells (CD45^+^ CD3^+^ PD-1^+^), T helper cells (CD45^+^ CD3^+^ CD4^+^), cytotoxic T cells (CD45^+^ CD3^+^ CD8^+^), T regulatory cells (CD45^+^ CD3^+^ CD4^+^ CD25^+^ FoxP3^+^), as well as the ratio CD8/Treg in the spleens at day 24 of the study (day 7 post-treatment). Values represent the mean ± SEM (n = 5 mice per group). Data were analyzed using an ANOVA with a Dunnett’s post hoc test, ^*^P < 0.05.

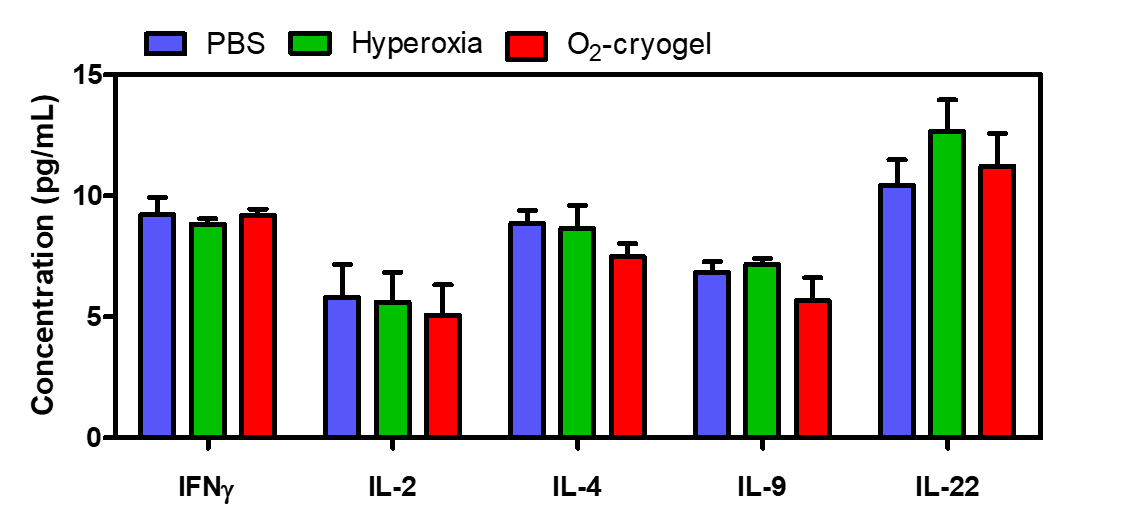

**Supplementary Figure 8. Serum cytokine levels following O_2_-cryogel co-adjuvant therapy.** Quantification of IFNγ, IL-2, IL-4, IL-9 and IL-22 cytokine levels in blood at day 7 post-treatment (day 24 of the study). Values represent the mean ± SEM (n = 5 mice per group). Data were analyzed using an ANOVA with a Dunnett’s post hoc test, ^*^P < 0.05.
